# The histidine phosphotransfer AHP4 plays a negative role in *Arabidopsis* plant response to drought

**DOI:** 10.1101/2020.07.30.229971

**Authors:** Chien Van Ha, Kien Huu Nguyen, Mohammad Golam Mostofa, Cuong Duy Tran, Yasuko Watanabe, Weiqiang Li, Yuriko Osakabe, Mayuko Sato, Kiminori Toyooka, Maho Tanaka, Motoaki Seki, David J. Burritt, Cheyenne Anderson, Ru Zhang, Lam-Son Phan Tran

## Abstract

Cytokinin plays an important role in plant stress responses via a multistep signaling pathway, involving the histidine phosphotransfer proteins (HPs). In *Arabidopsis thaliana*, the AHP2, AHP3 and AHP5 proteins are known to impact drought responses; however, the role of AHP4 in drought adaptation remains undetermined. In the present study, using a loss-of-function approach we showed that AHP4 possesses a negative regulatory role in *Arabidopsis’*s response to drought. This is evidenced by both higher survival rates of *ahp4* than wild-type (WT) plants under drought conditions, and the down-regulated *AHP4* expression in WT during periods of dehydration. Comparative transcriptome analysis of *ahp4* and WT plants revealed AHP4-mediated expression of several dehydration- and/or abscisic acid (ABA)-responsive genes involved in regulation of various physiological and biochemical processes important for plant drought acclimation. In comparison with WT, *ahp4* plants showed increased wax crystal accumulation in stems, thicker cuticles in leaves, greater sensitivity to exogenous ABA at germination, narrow stomatal apertures, heightened leaf temperatures during dehydration, and longer root length under osmotic stress. Additionally, *ahp4* plants showed greater photosynthetic efficiency, lower levels of reactive oxygen species (ROS), reduced electrolyte leakage and lipid peroxidation, and increased anthocyanin contents under drought, when compared with WT. These differences displayed in *ahp4* plants are likely due to up-regulation of genes that encode enzymes involved in ROS-scavenging and non-enzymatic antioxidant metabolism. The role of AHP4 in negative regulation of multiple protective mechanisms associated with drought tolerance could make editing of *AHP4* a promising approach for the production of drought-tolerant crop plants.

**Significance statement:** Loss-of-function analysis of the cytokinin signaling member AHP4 revealed its function in *Arabidopsis* adaptation to drought as a negative regulator, affecting various physiological and biochemical processes by modulating the expression of a large set of genes potentially in a crosstalk with ABA. *AHP4* and its homologs are promising candidates for gene editing to develop drought-tolerant crop cultivars.

## INTRODUCTION

Abiotic stresses, including drought, have detrimental effects on the growth and productivity of many important crops, resulting in significant yield losses that may lead to food shortages and threaten agricultural sustainability (Daryanto et al., 2016; Abdelrahman et al., 2018; Lamaoui et al., 2018). To cope with environmental stresses, plants have developed a range of cooperative physiological, biochemical and molecular mechanisms regulated by complex signaling networks (Sah et al., 2016; Choudhury et al., 2017). Phytohormones, the key regulators of plant growth and development, control different physiological and biochemical processes in plant responses to environmental stresses (Peleg and Blumwald, 2011; Verma et al., 2016; Mostofa et al., 2018; Wybouw and De Rybel, 2019). Among the various phytohormones, cytokinin (CK) is one of the central regulators in plant abiotic stress responses, and coordinates an array of functions, enabling plants to adapt to different types of stress (Ha et al., 2012; Zwack and Rashotte, 2015; Li et al., 2016; Cortleven et al., 2018; Li et al., 2019). CK has been reported to play a protective role in plant drought tolerance through enhancement of the endogenous CK level at the onset of drought in transgenic plants (Prerostova et al., 2018). More recently, by studying various CK-deficient and CK-signaling mutants, CK was found to play a negative regulatory role in plant drought responses (Li et al., 2016; Cortleven et al., 2018; Ramireddy et al., 2018). Thus, CK has been suggested to have multifaceted actions, playing both stimulatory and quiescent roles in plant drought tolerance (Prerostova et al., 2018).

The CK signaling pathway starts with histidine-kinase receptors (HKs), moves through histidine phosphotransfer proteins (HPs), and terminates in the activation of response regulators (RRs) which mediate the expression of downstream genes (Kieber and Schaller, 2018; Romanov et al., 2018). In *Arabidopsis thaliana*, there are three membrane-bound CK receptors (AHK2, AHK3 and CRE1/AHK4), which consist of a conserved CK-binding domain, a histidine kinase domain and a receiver domain (Keshishian and Rashotte, 2015; Pekarova et al., 2016; Romanov et al., 2018). Because the receptors are fixed to the membrane of endoplasmic reticulum, and RRs are primarily located in the nucleus, the intermediate HPs are necessary to help to relay the CK signal through the pathway (Keshishian and Rashotte, 2015; Romanov et al., 2018). There are five authentic AHPs (AHP1, 2, 3, 4 and 5) in *Arabidopsis*, which act as mediators in the multistep phosphorelay by transferring a phosphoryl group from the receiver domain of an activated AHK receptor to the receiver domain of an ARR (Keshishian and Rashotte, 2015; Kieber and Schaller, 2018). There are 24 RRs in *Arabidopsis* divided into two typical ARR groups, type-A (11 members) and type-B (10 members), and one atypical type-C (3 members) (Wybouw and De Rybel, 2019). Both type-A and type-B ARRs contain receiver domains; however, only type-B ARRs have a long C-terminal region that includes a MYB-like DNA-binding domain. Type-B ARRs, which play positive regulatory roles in CK signaling, function as transcription factors (TFs) in the final step of CK signaling by regulating downstream target gene expression, whereas type-A ARRs act as negative feedback regulators of CK signaling (Keshishian and Rashotte, 2015; Zwack and Rashotte, 2015; Kieber and Schaller, 2018; Romanov et al., 2018; Wybouw and De Rybel, 2019). Several type-B ARRs, namely ARR1, ARR2 and ARR12, were shown to be degraded by the kiss me deadly (KMD) F-box proteins (Kim et al., 2013).

Previous studies in *Arabidopsis* reported that several CK signaling members showed negative and redundant roles in drought responses of *Arabidopsis* plants, which included AHK2 and 3, AHP2, 3 and 5, type-B ARR1, 10 and 12 (Tran et al., 2007; Kang et al., 2012; Nishiyama et al., 2013; Kumar and Verslues, 2015; Nguyen et al., 2016). Among the 5 authentic AHPs, AHP4 is intriguingly distinct from the other AHPs not only evolutionarily, but also functionally. For instance, whereas AHP1, AHP2, AHP3, and AHP5 have been shown to act as positive regulators and have partially redundant functions in CK signaling, AHP4 was reported to play a negative regulatory role in CK signaling in some cases, such as later root formation (Hutchison et al., 2006). This raises a critical question regarding how the AHP4 acts within the CK signaling pathway in plant response to water deficit.

In the present study, using the loss-of-function approach, we initially characterize the detailed functions of AHP4 in regulating the response of *Arabidopsis* to drought. In contrast with its function in later root formation, our results indicate that AHP4 plays a negative role in plant response to drought similar to that played by AHP2, AHP3 and AHP5. Furthermore, this study showed increased responsiveness to ABA in *ahp4* mutant plants, which may contribute to higher drought tolerance of *ahp4* relative to wild-type (WT), and provide evidence for crosstalk between ABA and CK signaling. Additionally, higher leaf relative water content (RWC), lower leaf temperature, maintenance of higher cell membrane stability, increased reactive oxygen species (ROS)-scavenging enzyme activities and enhanced anthocyanin biosynthesis were all identified to potentially contribute to the enhanced drought tolerance of the *ahp4* mutant plants.

Comparative transcriptome analysis of the *ahp4* mutant and WT plants revealed several potential pathways, including ROS detoxification- and anthocyanin biosynthesis-related pathways, which were activated in *ahp4* plants under both normal and water-deficit conditions, further strengthening our findings.

## RESULTS

### *AHP4* is down-regulated in *Arabidopsis* plants during dehydration

To determine the involvement of *AHP4* in dehydration responses, we first used real-time quantitative PCR (RT-qPCR) to analyze the expression of the *AHP4* gene in *Arabidopsis* WT plants exposed to a dehydration treatment. Twenty-one-day-old WT plants were grown on plates of germination medium and exposed to dehydration treatment under ambient conditions. Rates of plant water loss were recorded during the treatment (Figure 1a). *AHP4* expression in both whole plants and shoots rapidly declined in response to dehydration (Figure 1b), with the lowest levels of expression of *AHP4* observed after 4 h of dehydration, followed by an incomplete recovery of expression after 6 h of dehydration (Figure 1b). *AHP4* expression in the roots of WT plants did not show the consistent responses observed in both whole plants and shoots under dehydration, but instead it showed a slight initial increase followed by a decline to a minimal value after 4 h of dehydration, finally followed by an increase in expression (Figure 1b). The well-known dehydration-responsive *responsive to desiccation 26* (*RD26*) gene (Fujita et al., 2004) was used as a marker to check the efficacy of the dehydration treatment (Figure 1c). The data presented in Figure 1c showed that the *RD26* expression levels in the shoots, roots and whole plants were elevated throughout the dehydration treatment, with the highest levels of expression found after 4 h of dehydration (∼76% water loss). Results of expression analysis suggest a dehydration-related function for *AHP4* in *Arabidopsis* plants.

**Figure 1.**
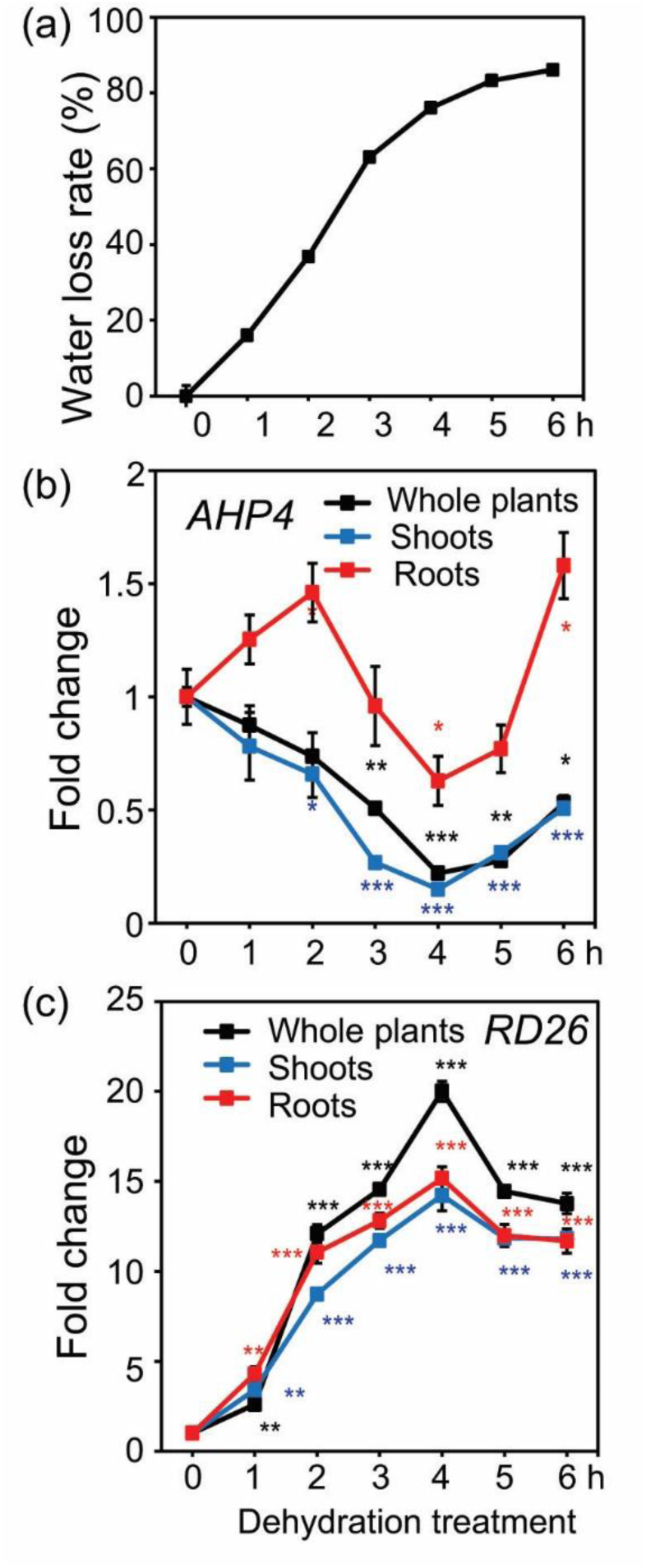
Expression patterns of the *AHP4* gene in *Arabidopsis* wild-type (WT) plants under dehydration treatment. (a) Water loss rate of 21-day-old WT plants grown on germination medium and subjected to a dehydration treatment. Data represent the means and standard errors (SEs) (*n* = 5). (b) Expression of the *AHP4* gene in 21-day-old WT plants subjected to dehydration treatment. (c) Expression of the stress-inducible *responsive to desiccation 26* (*RD26*) gene, which was used as a marker gene for checking the efficacy of dehydration treatment. Relative expression levels were normalized to a value of 1 in the respective control plants. Data represent the means and SEs (*n* = 3). Asterisks indicate significant differences as determined by a Student’s *t*-test (\**P* < 0.05, \*\**P* < 0.01, \*\*\**P* < 0.001).

### *Arabidopsis ahp4* mutant plants have improved drought tolerance

To further elucidate the role of the *AHP4* gene in plant response to drought, a loss-of-function approach was used. The *ahp4* mutant and WT plants were tested for drought tolerance using biomass reduction (Figure 2a-d) and survival assays (Figure 2e-k), and recovery (Figure 3). Under well-watered conditions, *ahp4* mutant plants showed insignificant difference in shoot growth compared with WT plants (Figure 2a-b, e, g). However, under water-deficit conditions *ahp4* mutant plants had a greater plant biomass than WT plants (Figure S2b). Compared with the well-watered plant groups, the stressed WT and *ahp4* mutant plants showed 37.1 and 17.7% biomass reduction, respectively (Figure S2c). Furthermore, *ahp4* plants had better recovery (Figure 3a-b) and higher survival rates than WT plants under water-deficit conditions (Figure 2f, h). Moreover, to test whether drought stress affects the photosynthetic system, we measured the maximum potential quantum efficiency of photosystem (PS) II (Fv/Fm), chlorophyll index and nonphotochemical chlorophyll fluorescence quenching (NPQ). Data showed that *ahp4* mutant plants exhibited greater Fv/Fm (Figure 3c-d), chlorophyll index (Figure 3e-f) and NPQ (Figure 3g-h) than WT plants under water-deficit conditions, suggesting that *ahp4* mutant plants had lower photoinhibition and higher photosynthetic efficiency under drought than WT plants. In addition, we observed that the primary roots of *ahp4* mutant plants grew better than that of WT plants, when they were treated with 200 or 300 mM of mannitol (Figure S1), suggesting that AHP4 controlled primary root growth in response to mannitol-induced water stress. Taken together, these results clearly indicated that loss-of-function of the *AHP4* gene resulted in a drought-tolerant phenotype, and that AHP4 acts as a negative regulator of responses involved in drought tolerance of *Arabidopsis* plants.

**Figure 2.**
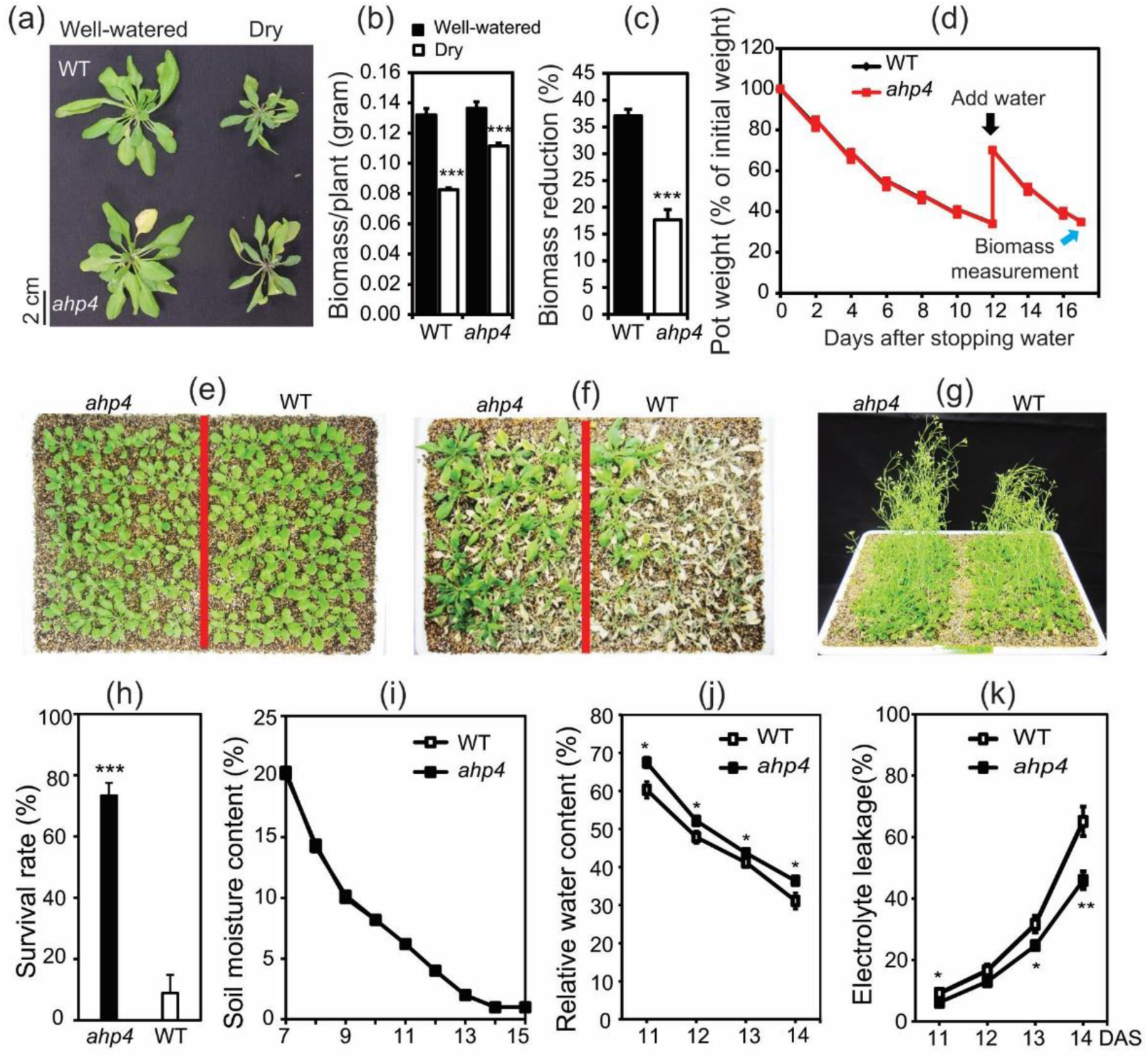
Drought-tolerant phenotype of the *ahp4* mutant plants. (a) Representative rosettes of *ahp4* and wild-type (WT) plants in the well-watered control and soil-drying treatments. (b) Biomass of *aph4* and WT plants under well-watered and soil-drying conditions. Data represent the means and standard errors (SEs) (*n* = 12/genotype). (c) Biomass reduction of soil-dried *aph4* and WT plants relative to respective well-watered control plants. Data represent the means and SEs (*n* = 12/genotype). (d) Averaged losses of *ahp4* and WT pot weights relative to initial pot weight during soil-drying (*n* = 12/genotype). Black arrow reveals when water was added to 70% of the initial pot weight. Blue arrow reveals when biomass was measured. (e) Two-week-old *ahp4* and WT plants were transferred from germination medium plates to soil and grown for one additional week. (f) Three-week-old plants were subjected to drought for 15 days and plants were photographed three days subsequent to rewatering and after removal of inflorescences. (g) Five-week-old plants were grown on the soil in well-watered control conditions. (h) Plant survival rates and SEs (*n* = 3, where each replicate represents the survival plant rate of 30 plants/genotype). (i) Soil moisture content was recorded during the water withholding (*n* = 5 positions/genotype/day). (j) Relative water contents of *ahp4* and WT plants grown and subjected to water withholding treatment as described in (e-f). Data represent the means and SEs (*n* = 5/genotype). (k) Electrolyte leakage rates of *ahp4* and WT plants grown and subjected to drought treatment as described in (e-f). Data represent the means and SEs (*n* = 5/genotype). Asterisks indicate significant differences between the two genotypes as determined by a Student’s *t*-test (\**P* < 0.05, \*\**P* < 0.01, \*\*\**P* < 0.001). DAS, days after stress.

**Figure 3.**
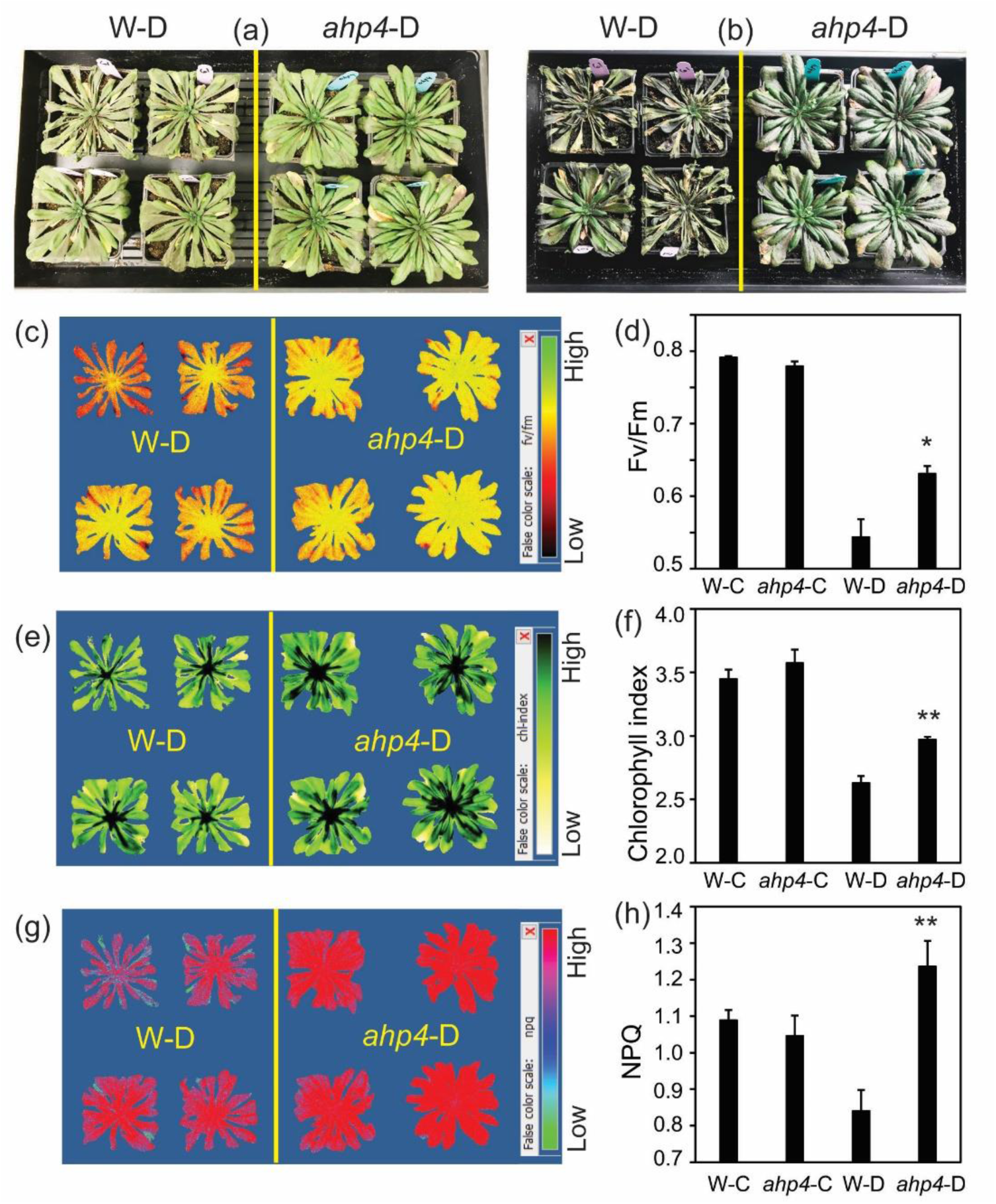
Drought-tolerant phenotype, maximum potential quantum efficiency of photosystem II (Fv/Fm), chlorophyll (Chl) index and nonphotochemical Chl fluorescence quenching (NPQ) of the *ahp4* mutant and wild-type (WT) plants. (a) Representative of *ahp4* and WT plants after 14 days of drought. (b) Representative of *ahp4* and WT plants were exposed to drought for 15 days, and then rewatered for 3 days. (c) Representative image of Fv/Fm of *ahp4* and WT plants after 14 days of drought. (d) The Fv/Fm of 49-day-old *ahp4* and WT plants under well-watered control conditions and after 14 days of drought treatment. (e) Representative image of Chl index of *ahp4* and WT plants after 14 days of drought. (f) Chl index of 49-day-old *ahp4* and WT plants under well-watered control conditions and after 14 days of drought treatment. (g) Representative image of NPQ of *ahp4* and WT plants at 14 days of drought. (h) NPQ of 49-day-old *ahp4* and WT plants under well-watered control conditions and after 14 days of drought treatment. Data represent the means and standard errors (SE) (*n* = 4). Asterisks indicate significant differences as determined by a Student’s *t*-test (\**P* < 0.05, \*\**P* < 0.01). W-C, WT under well-watered control; W-D, WT under drought; *ahp4*-C, *ahp4* under well-watered control; *ahp4*-D, *ahp4* under drought.

### Comparative transcriptome analyses of *ahp4* and WT plants under non-stressed and dehydration conditions

Based on *AHP4* expression and drought-tolerant test results, we performed a microarray analysis to investigate AHP4-dependent CK signaling-mediated downstream genes involved in plant adaptation to water deficiency. The experimental design for comparing transcriptome data obtained from the leaves of *ahp4* and WT plants under non-stressed and dehydration conditions are illustrated in Figure S2a, and the results of the microarray analysis are summarized in Figure S2b-d and Table S1. Confirmation of the microarray analysis, which was carried out using RT-qPCR of five selected genes, supported the results of microarray data (Figure S3). Compared with WT plants under non-stressed conditions (M-C/W-C), the analysis of differentially expressed genes (DEGs) found 1544 up-regulated and 753 down-regulated genes (Figure S2c; Table S2a-b). Comparison of DEGs in *ahp4* versus WT plants under dehydration (M-D/W-D) revealed a total of 1432 up-regulated genes (Figure S2c; Table S3a), with more genes being up-regulated after 2-h (M-D2/W-D2) dehydration than after 5-h (M-D5/W-D5) dehydration (Table S3b-c). A smaller number of genes (973) were down-regulated in (M-D/W-D) (Figure S2c; Table S3d), with more genes being repressed with increased duration of dehydration (Table S3e-f).

Using Venn diagram analysis, many dehydration-inducible genes belonging to the groups of genes were identified as being up-regulated in the M-C/W-C and/or M-D/W-D comparisons (Figure S2d, *i*-*ii*; Tables S2a, S3a and S4a-b). Dehydration-repressible genes were also found to belong to the groups of genes that were down-regulated genes in the M-C/W-C and/or M-D/W-D comparisons (Figure S2d, *iii*-*iv*; Tables S2b, S3d and S4c-d). These changes in gene expression likely resulted in drought-tolerant enhancement in *ahp4* mutant plants. MapMan analysis was then used to classify the DEGs, identified from the M-C/W-C and/or M-D/W-D comparisons, into functional groups (Figure S4), and to provide an overview of changes in general metabolism in *ahp4* versus WT plants (Figure S5) under non-stressed and dehydration conditions.

A detailed analysis of the DEGs obtained from the comparison of *ahp4* versus WT transcriptomes under non-stressed and dehydration conditions was then conducted to identify dehydration- and/or ABA-inducible genes, whose up-regulation might contribute to the drought-tolerance of the *ahp4* mutant plants. A number of dehydration- and/or ABA-inducible DEGs were annotated to encode AP2- and MYB-type TFs, C2H2-like zinc finger proteins, leucine-rich repeat kinases, glycine-aspartate-serine-leucine (GDSL) esterase/lipase, late embryogenesis abundant (LEA) proteins, α/β-hydrolases, β-glucosidases and glycoside hydrolases (Tables S2 and S3). Many of the dehydration-inducible genes found to be up-regulated in dehydrated *ahp4* versus dehydrated WT plants (M-D/W-D) were associated with secondary metabolism, which included genes that are known to be involved in flavonoid biosynthesis like those encoding flavonol synthase 4, UDP-glucosyl transferase 73C6 and UDP-glucosyl transferase 78D1 (Figures S5 and S6, Table S3a). Furthermore, several genes associated with cuticular wax biosynthesis, including *AT1G34490, AT1G34500, AT5G51420* and *AT5G55360*, were found to be up-regulated in *ahp4* mutant plants under dehydration (M-D/W-D) (Figures S5 and S6, Table S3a). In addition, several ROS-related genes were found to have altered expression patterns in dehydrated *ahp4* versus dehydrated WT plants, suggesting that alteration in ROS metabolism could also participate in drought tolerance of *ahp4* mutant plants (Figure S4, Tables S2 and S3). In summary, the above results suggest that the enhanced drought tolerance of *ahp4* mutant plants (Figures 2-3) was the result of differential regulations of a set of DEGs associated with a range of metabolic processes.

### Reduced stomatal apertures, induced photosynthetic efficiency and enhanced ABA responsiveness in *ahp4* plants

Plants respond to drought/dehydration stress by inducing stomatal closure and/or reducing stomatal density which help them retain water under water-deficit conditions. To determine the role of AHP4 in controlling stomatal activity, we conducted several assays under both normal and dehydration conditions. As reported earlier under drought (Figure 2j) and long-term dehydration (Figure S2a) conditions, *ahp4* plants also showed greater RWC than WT plants during the short-term dehydration (Figure 4a). Furthermore, *ahp4* plants exhibited higher leaf temperatures than WT plants under both non-stressed and the same dehydration conditions (Figure 4b), suggesting a relationship between water transpiration and stomatal density and/or stomatal movement. Because no significant differences were observed in stomatal density between *ahp4* mutant and WT plants (Figure 4c), the higher leaf temperatures observed in *ahp4* plants (Figure 4b) resulted from their narrower stomatal apertures in comparison with WT plants under both non-dehydrated and dehydrated conditions as shown in Figure 4d-e. Additionally, using different light intensity treatments, we found that the stomatal conductance and transpiration rates of *ahp4* mutant showed lower trends than WT plants under non-stressed conditions, but they were comparable under dehydration conditions (Figure S7).

**Figure 4.**
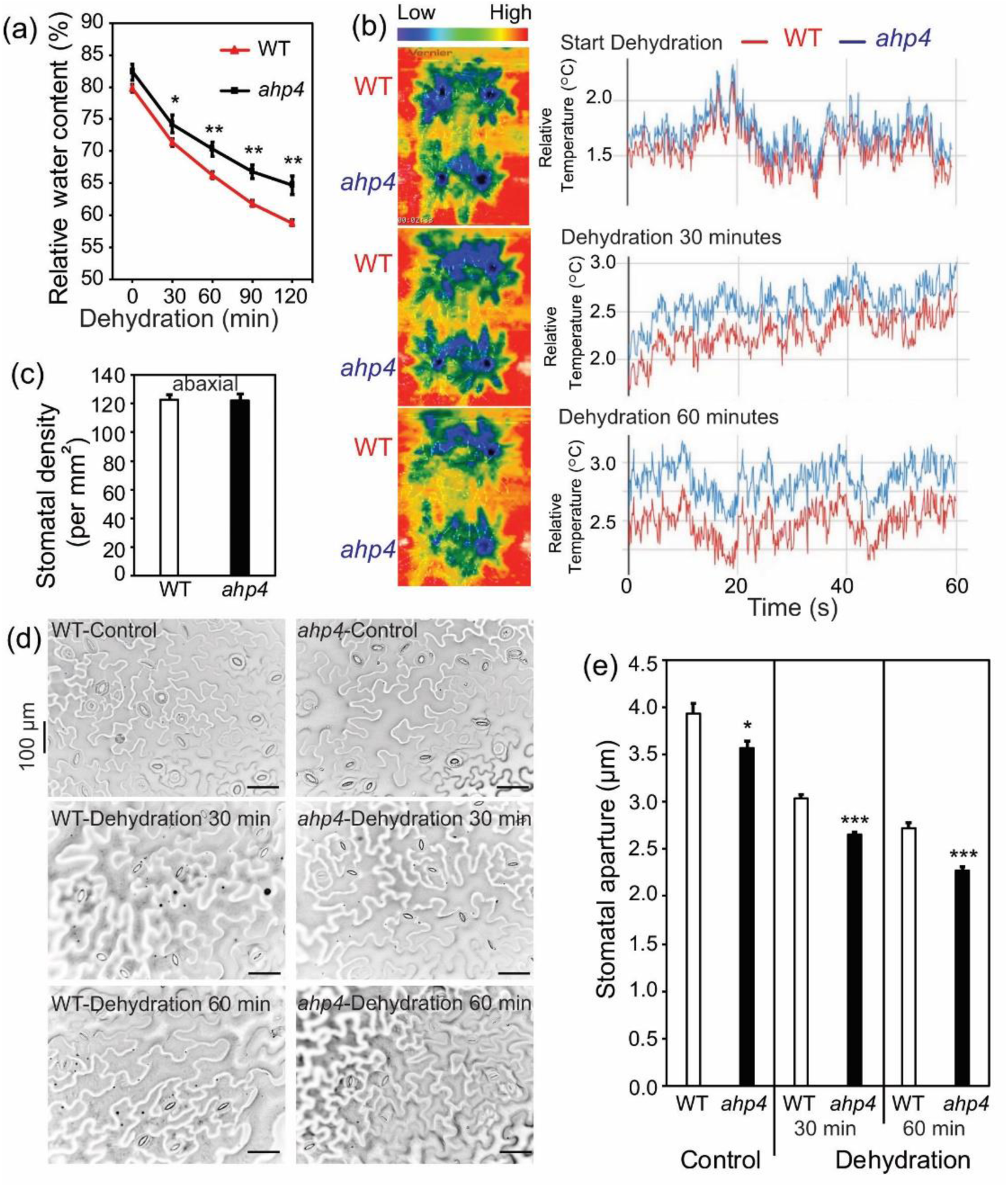
Relative water content (RWC), relative leaf temperatures, stomatal densities and stomatal apertures of *ahp4* mutant and wild-type (WT) plants. (a) RWC of 32-day-old soil-grown *ahp4* and WT plants subjected to a dehydration treatment. Data represent the means and standard errors (SEs) (*n* = 5). (b) Relative leaf temperatures of 32-day-old soil-grown *ahp4* and WT plants were recorded during the dehydration treatment. The rainbow color scale indicates the relative temperatures. (c) Average stomatal densities of rosette leaves from 32-day-old soil-grown *ahp4* and WT plants. Stomata were counted from eight different areas on each leaf. Data represent the means and SEs (*n* = 5). (d) Guard cells of 32-day-old soil-grown *ahp4* and WT plants were subjected to a dehydration treatment for 0 (Control), 30 and 60 minutes. (e) Average size of the stomatal aperture of rosette leaves from 32-day-old soil-grown *ahp4* and WT plants subjected to a dehydration treatment for 0 (Control), 30 and 60 minutes. Data represent the mean and SEs (*n* = 5 plants/genotype; for each plant the average of 22 stomatal measurements from a single leaf was calculated). Asterisks indicate significant differences between the two genotypes as determined by a Student’s *t*-test (*\*P* < 0.05, \*\**P* < 0.01, \*\*\**P* < 0.001).

In addition, ABA-mediated regulation of stomatal opening/closure, and/or ABA-associated mechanisms that promote cellular dehydration tolerance are all important for drought tolerance in plants (Osakabe et al., 2014; Kuromori et al., 2018). To determine if any such ABA-related relationships could be responsible for the differences in the RWC of *ahp4* and WT plants and contributed to the differences in drought tolerance, we next compared the stomatal apertures of the two genotypes with or without exogenous ABA treatment. In agreement with stomatal closure data shown in Figure 4d-e under non-stressed conditions, *ahp4* plants showed narrower stomatal apertures than WT did in the absence of ABA; however, both *ahp4* and WT showed comparable stomatal apertures in the presence of ABA (Figure S8). This collective data demonstrated that the lower rate of water loss in *ahp4* mutant plants, relative to that of WT plants, was caused by a decrease in water transpiration rate due to the impairment of stomatal closure, contributing to the increased drought tolerance of *ahp4* mutant plants.

To determine if increased drought tolerance of the *ahp4* mutant plants was also associated with increased ABA responsiveness, which may induce downstream ABA-responsive genes that are not related to stomatal activity processes (Fujita et al., 2005), we conducted a seed germination and root growth assay on medium supplemented with different concentrations of ABA. Results showed that the *ahp4* mutant plants were more sensitive to ABA than WT plants at both germination and seedling stages (Figure S9a-d). Furthermore, we observed that the expression of *AHP4* gene in WT plants was significantly down-regulated in whole plants and roots treated with ABA at all the time points of dehydration period (Figure S9e). Interestingly, *AHP4* expression was mostly un-changed in shoots of ABA-treated WT plants at earlier time points, and was then significantly down-regulated after 6 h of dehydration (Figure S9e). These findings indicate that enhanced response of *ahp4* mutant plants to ABA might contribute to their improved drought tolerance, and repression of *AHP4* transcription by stress-induced ABA might help plants adapt to adverse environmental conditions.

### Enhancement of cell membrane integrity and cuticle thickness of *ahp4* plants

The greater RWC observed in *ahp4* mutant plants compared with WT plants under water deficit suggests the possibility of changes in cell membrane integrity (Figures 2j, 4a and S2a) (Verslues et al., 2006). To determine if there were any differences in cell membrane integrity induced by water deficit, tissue electrolyte leakage in *ahp4* and WT plants under drought was measured (Figure 2k). Our data demonstrated that the loss-of-function mutation in *AHP4* led to a considerably lower electrolyte leakage in *ahp4* mutant than in WT plants under water deficit (Figure 2k).

The cuticle, which composes of cutin and waxes, covers the epidermis and controls the water permeability, limiting non-stomatal water loss (Sieber et al., 2000; Yeats and Rose, 2013; Jetter and Riederer, 2016). We next evaluated whether there is difference in cuticle structure between *ahp4* and WT plants, which would differentiate the transpiration rates of *ahp4* mutant and WT plants. First, we conducted a chlorophyll leaching assay of rosette leaves of *ahp4* mutant and WT plants grown under normal conditions to compare cuticle permeability. The chlorophylls were found to leach more slowly from *ahp4* leaves than WT leaves (Figure 5a), suggesting lower cuticle permeability in *ahp4* compared with WT plants. Second, toluidine blue (TB) staining images showed lower TB uptake by the leaves of *ahp4* mutant as compared with that of WT plants (Figure 5b). These data infer that loss-of-function of *AHP4* might result in enhanced cuticle structure of *ahp4* mutant plants, which could prevent non-stomatal water loss under adverse conditions. Third, scanning electron microscopy (SEM) imaging of the surface wax ornamentation of the stems of *ahp4* and WT plants indicated a higher density of surface wax in *ahp4* mutant versus WT plants (Figure 5c). Fourth, transmission electron microscopy (TEM) imaging of the cuticle layers of the fifth leaves showed thicker cuticle layer in *ahp4* mutant versus WT plants (Figure 5d-e). Taken together, these results clearly indicate that increased cuticle thickness in *ahp4* mutant plants may help to prevent non-stomatal water loss from leaf and stem surfaces. To investigate the underlying genetics associated with increased cuticle thickness, transcript levels of several genes involved in wax biosynthesis, namely *decrease wax biosynthesis* (*DEWAX*), *fatty acyl-coenzyme A reductase 6* (*FAR6*) and *shine 1* (*SHIN1*), were compared in *ahp4* mutant and WT plants (Cui et al., 2016). Results revealed that the expression of *DEWAX* was down-regulated in *ahp4* mutant under dehydration conditions, while that of *FAR6* and *SHIN1* was up-regulated in *ahp4* mutant plants under both non-dehydrated and dehydrated conditions (Figure 5f). This difference in expression levels might contribute to increased cuticle thickness (Figure 5d-e). These collective results suggest that AHP4 may play an important role in controlling wax biosynthesis in *Arabidopsis* plants acclimatizing to water stress.

**Figure 5.**
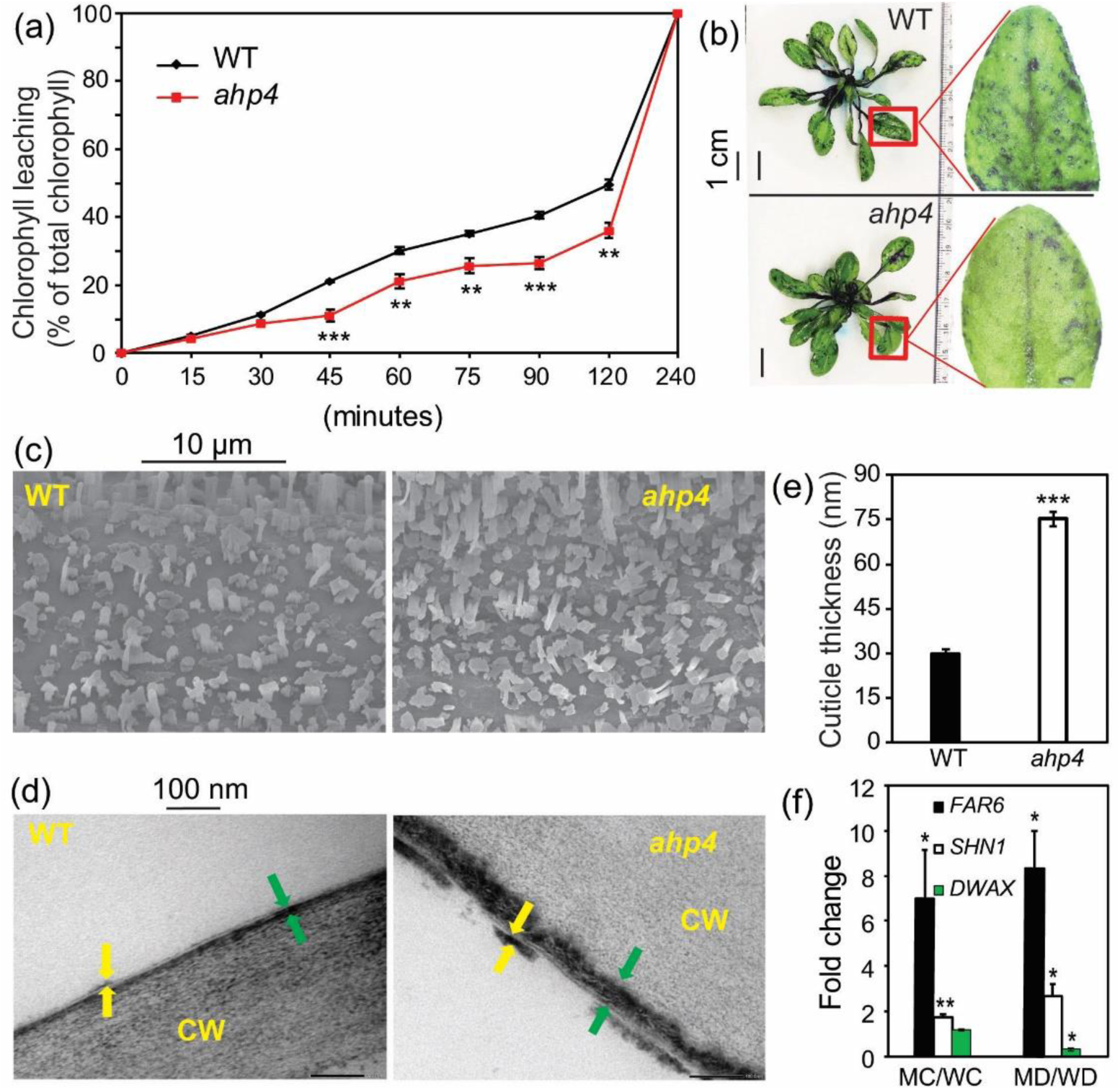
Chlorophyll leaching and toluidine blue (TB) staining patterns, wax accumulation, cuticle thickness and wax-related gene expression of *ahp4* mutant and wild-type (WT) plants. (a) Fourteen-day-old *ahp4* and WT plants grown on germination medium plates were transferred to soil and grown under well-watered conditions for 21 additional days. Chlorophyll leaching of 35-day-old *ahp4* and WT plants were measured at indicated time points. Data represent the means and standard errors (*n* = 5). (b) TB staining patterns of leaves of 35-day-old *ahp4* and WT plants grown on soil as described in (a). (c) Wax surface ornamentation of stems of 35-day-old *ahp4* and WT plants grown on soil as described in (a) were detected by scanning electron microscope. (d) Cuticle of the fifth leaves (adaxial side) of 35-day-old *ahp4* and WT plants grown on soil as described in (a) were also observed by transmission electron microscope. The green arrows indicate cuticular layer, and yellow arrows indicate cuticle proper. (e) Cuticle thickness of the fifth leaves (adaxial side) of *ahp4* and WT plants was measured by ImageJ software. Data represent the means and SEs (*n* = 5, where each repeat was counted from eight different areas). (f) Expression of several wax-related genes in *ahp4* and WT plants with or without 5 h of dehydration treatment as described in **Fig. S2**. Data represent the means and SEs (*n* = 3). Asterisks indicate significant differences between the two genotypes as determined by a Student’s *t*-test (\**P* < 0.05, \*\**P* < 0.01, \*\*\**P* < 0.001). CW, cell wall; MC/WC, *ahp4* well-watered control 0 h versus WT well-watered control 0 h; MD/WD, *ahp4* dehydrated 5 h versus WT dehydrated 5 h. *DEWAX, decrease wax biosynthesis*; *FAR6, fatty acyl-coenzyme A reductase 6*; *SHIN1, shine 1*.

### Decreased ROS accumulation and oxidative damage, and increased ROS-scavenging antioxidant capacity in *ahp4* plants

Plants respond to various environmental stresses, including drought, via many adaptive mechanisms, including maintaining a balance between ROS production and detoxification, which is important for the maintenance of many cellular functions (Wang et al., 2015; Choudhury et al., 2017; Farooq et al., 2019; Huang et al., 2019; Xie et al., 2019). As the microarray analyses showed that AHP4 is involved in regulating the expression of genes involved in ROS metabolism, we decided to further investigate the role of AHP4 in ROS homeostasis. We did so by examining ROS production (O_2_^•−^ and H_2_O_2_) in the leaves of *ahp4* mutant and WT plants under well-watered and water-deficit conditions using the staining approach (Figure 6a-b). We observed a comparable accumulation of O_2_^•−^ and H_2_O_2_ in the leaves of well-watered *ahp4* and WT plants. In contrast, under water deficit conditions, the levels of O_2_^•−^ and H_2_O_2_ in *ahp4* mutant remained much lower than in WT plants (Figure 6a-b). To determine if oxidative damage was in fact reduced in *ahp4* mutant, the contents of malondialdehyde (MDA), a byproduct of lipid peroxidation and common marker of oxidative stress (Mostofa et al., 2015), in *ahp4* and WT plants were compared. While no significant difference in MDA content was observed in *ahp4* mutant and WT plants under well-watered conditions, after 13 days of drought stress *ahp4* mutant displayed much lower MDA level than did the WT (Figure 6c). These data suggest that loss-of-function of *AHP4* mitigates oxidative damage in *Arabidopsis* plants by reducing drought-induced ROS accumulation and lipid peroxidation.

**Figure 6.**
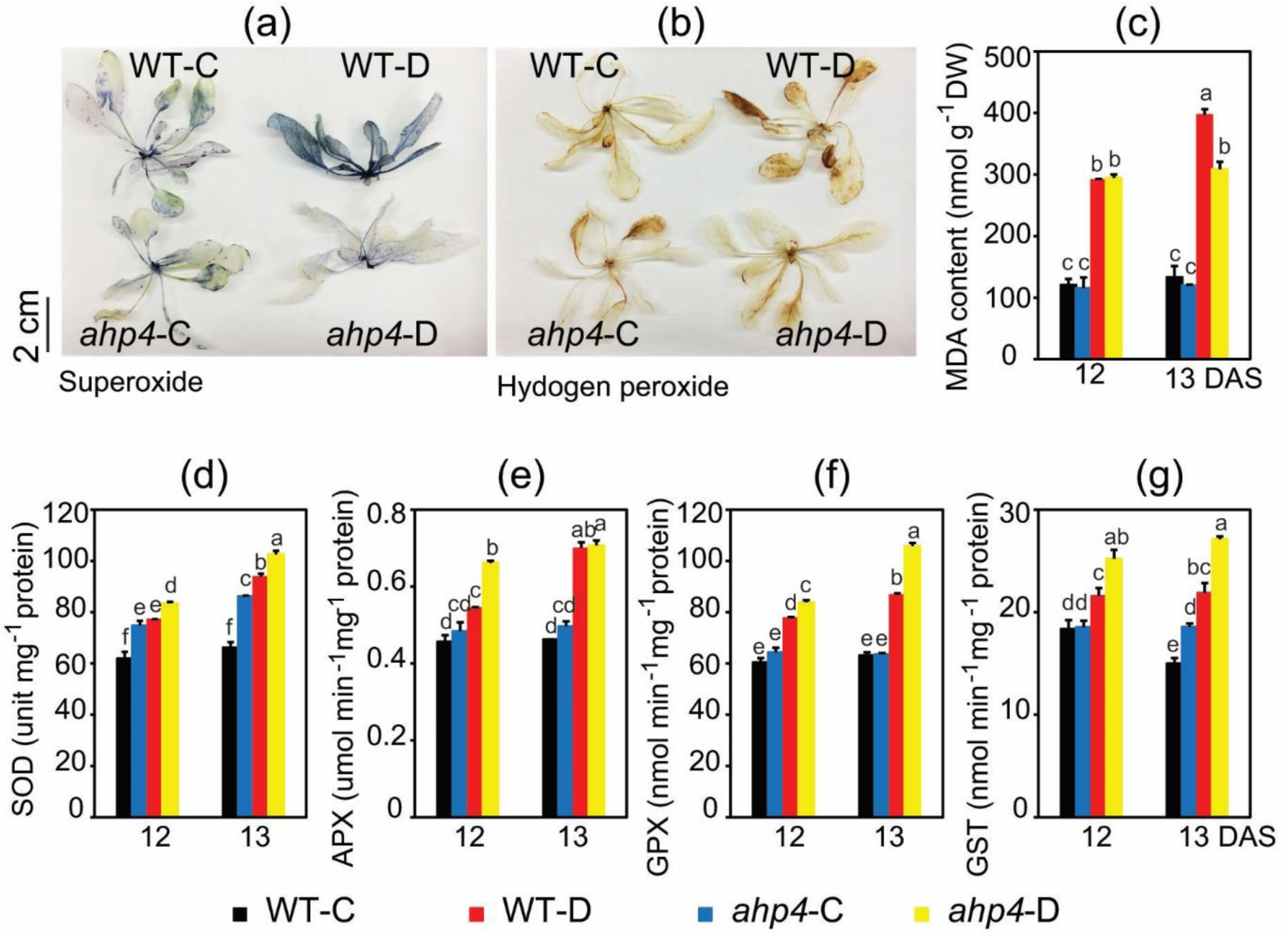
Determination of reactive oxygen species accumulation, malondialdehyde (MDA) content and antioxidant enzyme activities in *ahp4* mutant and wild-type (WT) plants during drought. (a) Nitro blue tetrazolium staining of superoxide; and (b) diamino-benzidine staining of hydrogen peroxide in rosettes of *ahp4* and WT plants exposed to drought for 12 days. (c) MDA content of *ahp4* and WT plants under drought. Data represent the means and standard errors (SEs) (*n* = 3). (d) Superoxide dismutase (SOD); (e) ascorbate peroxidase (APX); (f) glutathione peroxidase (GPX); and (g) glutathione S-transferase (GST) activities in *ahp4* and WT plants under soil-drying. Data represent the means and SEs (*n* = 3). Different superscripted letters (a, b, c, d, e and f) within the column reveal statistically significant differences between the two genotypes, and among the treatments, which were determined by Duncan’s multiple range test (*P* < 0.05). DAS, days after stress; WT-C, wild-type control; WT-D, wild-type drought; *ahp4*-C, *ahp4* control; *ahp4*-D, *ahp4* drought.

The results on drought-induced oxidative stress combined with the transcriptome data indicate that under the water-deficit conditions *ahp4* mutant plants have a greater antioxidant capacity to more efficiently detoxify ROS, when compared with WT plants. To investigate this further, first the activities of several key ROS-scavenging enzymes like superoxide dismutase (SOD), ascorbate peroxidase (APX), glutathione peroxidase (GPX) and glutathione S-transferase (GST) were determined in WT and *ahp4* plants under well-watered and water-deficit conditions (Figure 6d-g). Under well-watered conditions, no differences were observed in the activities of APX, GPX and GST in the leaves of WT and *ahp4* mutant plants, and only SOD activity showed higher activity in *ahp4* mutants than in WT plants. In contrast, under water-deficit conditions the activities of the examined ROS-scavenging enzymes were generally greater in the leaves of *ahp4* than in WT plants (Figure 6d-g). Second, although there were no significant differences in anthocyanin contents between *ahp4* mutant and WT plants under well-watered conditions (Figure S10a-b), under water-deficit conditions *ahp4* mutant plants possessed higher anthocyanin contents than WT plants (Figure S10a-b). Concurrent with this observation, under water-deficit conditions, the expression of genes involved in anthocyanin/flavonoid biosynthesis showed a greater up-regulation in *ahp4* mutant than in WT plants (Figure S10c). These findings suggest that enhanced accumulation of anthocyanins, which may act as an antioxidant under different types of environmental stress, including drought (Nakabayashi et al., 2014; Lotkowska et al., 2015; Nguyen et al., 2016; Nguyen et al., 2016; Li et al., 2017), was associated with the drought-tolerant phenotype of *ahp4* mutant plants. Collectively, these results suggested that loss-of function of AHP4 enhanced antioxidant defense as a preventive measure to protect *ahp4* plants from drought-induced oxidative damage.

## DISCUSSION

Globally abiotic stresses, including drought, are the primary factors that impact plant survival and cause yield losses in crop plants (Lamaoui et al., 2018). To acclimatize to environmental stresses, plants need to be able to regulate a variety of developmental, physiological and cellular mechanisms to survive (Osakabe et al., 2014; Sah et al., 2016; Choudhury et al., 2017; Zandalinas et al., 2018). In this study, we found that the expression of *AHP4* gene was down-regulated by dehydration or ABA treatment (Figure 1b and S9e), suggesting a possible involvement of *APH4* in negative regulation of *Arabidopsis* adaptation to drought in ABA-dependent manner. This hypothesis is strongly supported by the fact that the *aph4* mutant plants exhibited a drought-tolerant phenotype (Figures 2-3). Next, we used various physiological and biochemical assays and a genome-wide transcriptome analysis to investigate, identify and critically evaluate numerous AHP4-regulated genes, several developmental and physiological mechanisms, and selected pathways that might contribute to AHP4-mediated drought responses in *Arabidopsis*. Our results demonstrate that AHP4 does indeed play a negative regulatory role in a wide range of mechanisms, as summarized in Figure 7.

**Figure 7.**
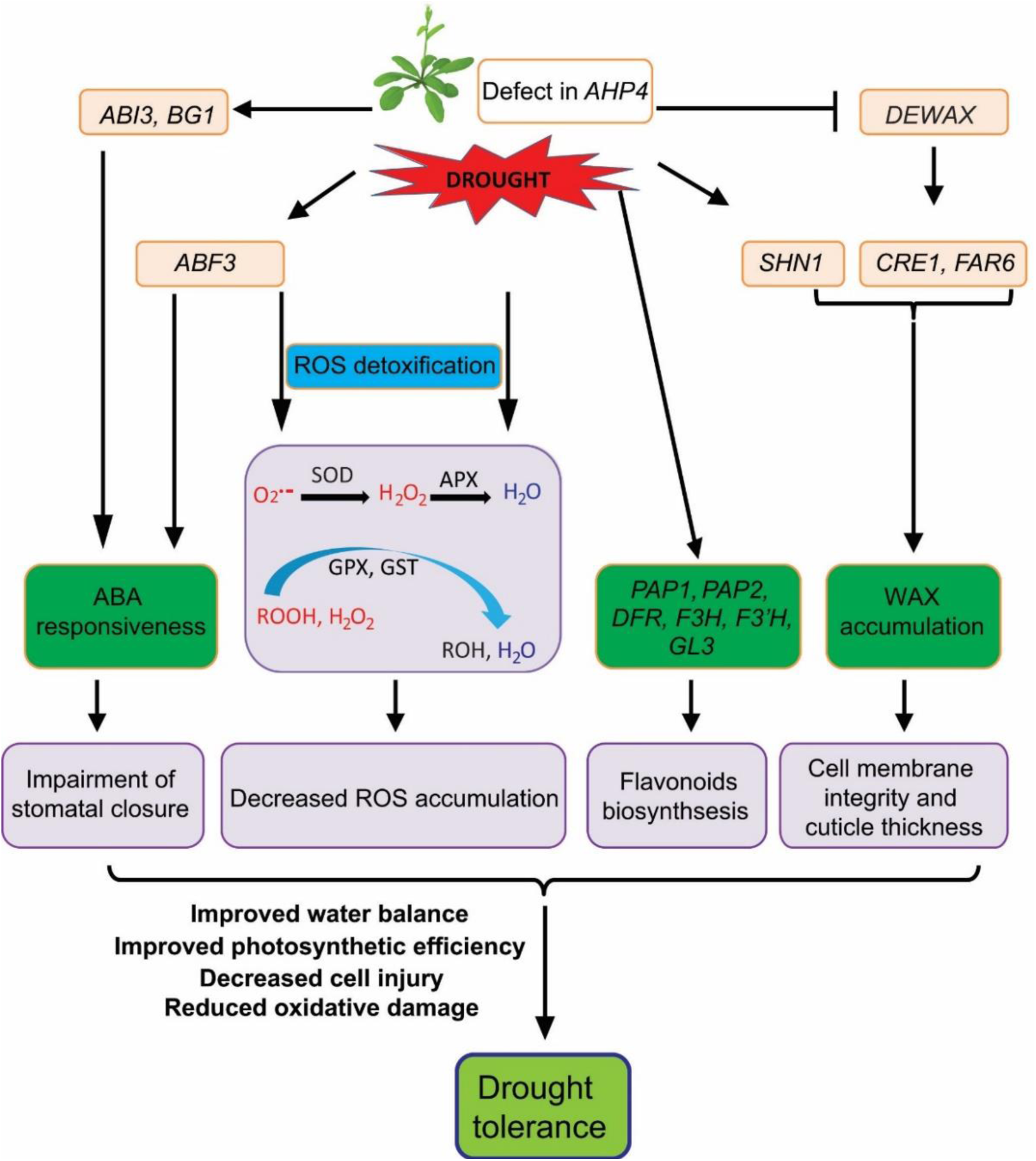
Model for negative regulatory role of AHP4 in response of *Arabidopsis thaliana* to drought. Under drought, expression of *AHP4* is repressed, weakening the action of the cytokinin signaling. Downregulation of *AHP4* results in changes in various physiological and biochemical processes, including impairment of stomatal closure, enhanced photosynthetic efficiency, increased cell membrane integrity and cuticle thickness, and improvement of reactive oxygen species (ROS) detoxification through increasing the levels of both enzymatic and non-enzymatic antioxidants. ABA, abscisic acid; *ABF3, ABA-responsive element-binding factor 3*; *ABI3, ABA insensitive 3*; APX, ascorbate peroxidase; *BG1, β-glucosidase 1*; *DEWAX, decrease wax biosynthesis*; *DFR, dihydroflavonol 4-reductase*; *F3H, flavanone 3-hydroxylase*; *F3*^*’*^*H, flavonoid 3*^*’*^*-monooxygenase*; *FAR6, fatty acyl-coenzyme A reductase 6*; *GL3, glabra 3*; GPX, glutathione peroxidase; GST, glutathione S-transferase; ROOH, organic hydroperoxides; ROH, organic hydroxyl; *PAP1/PAP2, production of anthocyanin pigment 1/2*; *SHIN1, shine 1*; SOD, superoxide dismutase.

First, we investigated if loss-of-function of *AHP4* would enhance primary root growth in plants exposed to concentrations of mannitol high enough to cause osmotic stress. Around 200 or 300 mM mannitol, *ahp4* mutant plants showed increased primary root growth compared with WT plants (Figure S1). These results suggest that the enhancement of primary root growth observed in *ahp4* mutant plants under osmotic stress treatment might contribute to *ahp4* mutant’s drought-tolerant phenotype. Increasing downward root growth is an essential mechanism which can enable plants to absorb water from deep layers within the soil when water in shallower soil layers is limited (Feng et al., 2016). The negative regulatory role of *AHP4*, and thus CK signaling, in root growth and development observed in this study under osmotic stress, is supported by previous studies which reported significant increases in root length and biomass in both transgenic plants with reduced CK content (Werner et al., 2010) and CK-signaling *ahk2,3* double mutant (Riefler et al., 2006; Li et al., 2019) even under normal growth conditions, and enhanced root elongation in *ahk3* mutant plants exposed to low water potential (Kumar and Verslues, 2015). Additionally, the CK-deficient and *ahk2,3* plants were better able to survive under drought (Tran et al., 2007; Werner et al., 2010; Nishiyama et al., 2011; Kang et al., 2012).

Second, the drought tolerance of *ahp4* mutant plants was found be enhanced by their ability to retain higher RWC and Fv/Fm when compared with WT (Figures 2j, 3c-d, 4a and S2a) to better survive water deficit (Figures 2-3). Maintaining higher RWC and better drought tolerance requires the down-regulation of *APH4* in order to activate a range of biochemical and physiological mechanisms that help plants cope with water deficit as visualized in the model shown in Figure 7. These mechanisms include increased maximum photochemical efficiency of PSII (Figure 3c-d), reduced stomatal apertures (Figures 4d-e and S8), decreased cell membrane damage and cuticle permeability (Figures 2k and 5), increased ABA responsiveness (Figure S9), and increased antioxidant defense (Figures 6 and S10).

During drought, plants alter the photosynthetic machinery to maintain cell water status for better survival (Wang et al., 2018). For evaluation of drought tolerance of plants in terms of their ability of maintain photosynthesis, the Fv/Fm, stomatal conductance and transpiration rate parameters are widely used as photosynthetic indicators (Wang et al., 2018; Utsumi et al., 2019). Previous studies reported that the drought-tolerant plants have high Fv/Fm, NPQ, and low stomatal conductance and transpiration rate to maintain water status in plants during water scarcity (Wang et al., 2018; Utsumi et al., 2019). In this study, *ahp4* plants showed greater Fv/Fm and NPQ under drought (Figure 3c-d, g-h), higher leaf temperature during dehydration (Figure 4b), lower stomatal conductance and transpiration rates under well-watered conditions (Figure S7a-b) when compared with those in WT, suggesting that the enhanced drought tolerance of *ahp4* plants might be attributed to an alteration in photosynthetic responses. In addition, the observed greater photosynthetic efficiency of *ahp4* plants (Figure 3c-d) suggests that *ahp4* plants may absorb light better than WT plants. In support of this premise, the transcript levels of the photosystem light harvesting complex (LHC)-related genes, such as *LHCB2*.*1, 2*.*2, 2*.*3* and *4*.*1*, were up-regulated in *ahp4* plants when compared with WT plants under dehydration conditions (Table S5). These findings indicate the important role of AHP4 in regulation of photosynthetic processes in plant response to drought.

ABA is known to be a key factor in plant abiotic stress responses as it is involved in regulating the expression of many ABA- and/or stress-inducible genes (Osakabe et al., 2014; Osakabe et al., 2014; Ding et al., 2015; Sah et al., 2016). Previous studies demonstrated that ABA and CK signaling pathways have antagonistic actions in various processes, including germination, senescence and drought tolerance (Nishiyama et al., 2011; Wang et al., 2011; Nguyen et al., 2016; Abdelrahman et al., 2017; Huang et al., 2018). In the present study, the enhancement of the ABA response in *ahp4* mutant plants (Figure S9) could result in the up-regulation of downstream ABA- and/or stress-inducible genes under water deficit. These DEGs include the *LEA, ABA insensitive 3* (*ABI3*), *ABA-responsive element-binding factor 3* (*ABF3*) and *β-glucosidase 1* (*BG1*) genes (Table S3a). These genes are known to have various important functions, including membrane protection, maintenance of osmotic homeostasis and repair of cell damage, in plants under water deficit, leading to improvement of the overall performance of plants exposed to drought (Fujita et al., 2005; Verslues et al., 2006). *ABI3* encodes a AP2/ERF-type TF important for normal embryogenesis and seed development (Rohde et al., 2000; Roscoe et al., 2015), as well as for plants under dehydration stress (Bedi et al., 2016). ABI3 might have a positive regulatory function in *Arabidopsis* plants under drought as its transcription is up-regulated following dehydration, and its transcript levels are maintained during the subsequent stress recovery phases (Bedi et al., 2016; Bedi and Chaudhuri, 2018). On the other hand, the drought-induced *ABF3* encodes a member of the ABF*/*AREB subfamily of bZIP-type TFs (Fujita et al., 2013; Yoshida et al., 2015; Zandkarimi et al., 2015; Wang et al., 2016).

Overexpression/ectopic expression of *AtABF3* in many plant species, including *Arabidopsis*, rice (*Oryza sativa*), grape (*Vitis vinifera*), lettuce (*Lactuca sativa*), peanut (*Agrostis mongolica*), alfalfa (*Medicago sativa*) and cotton (*Gossypium hirsutum*), results in increased tolerance of transgenic plants to various stresses, including dehydration, cold, high temperatures and oxidative stress (Kang et al., 2002; Kim et al., 2004; E. Vanjildorj, 2005; Oh et al., 2005; Abdeen et al., 2010; Pruthvi et al., 2014; Wang et al., 2016; Kerr et al., 2017). These reports together strengthen the idea that up-regulation of *ABI3* and *ABF3* genes by loss-of-function of *AHP4* might contribute to enhanced drought tolerance of the *ahp4* mutant plants. Additionally, in the present study, the *AtBG1* gene, which encodes a β-glucosidase, was also expressed at higher levels in *ahp4* mutant plants than WT plants under water deficit (Table S3a). β-glucosidases release glucose from glucose-conjugates, such as the ABA-glucose ester by hydrolysis, releasing free ABA and increasing active ABA levels in plants (Lee et al., 2006; Dong et al., 2014). *BG1* expression is highly induced in WT plants treated with exogenous ABA, NaCl and high concentrations of mannitol (Dong et al., 2014). Overexpression of *AtBG1* in creeping bentgrass plants (*Agrostis stolonifera*) resulted in greater free ABA levels and increased drought tolerance in comparison with WT plants (Han et al., 2012). In contrast, *atbg1* mutant plants were shown to be more sensitive to drought than WT (Lee et al., 2006). The above findings support a role for AHP4, as a negative regulator of *AtBG1* expression, in the rapid ABA production pathway that is important for plants adapting to drought.

Under water deficit, plants commonly regulate water loss by controlling stomatal number and size of stomatal aperture (e.g. stomatal opening and closing) (Nilson and Assmann, 2007; Kerr et al., 2017; Saradadevi et al., 2017; Zandalinas et al., 2018). In the present study, a lower rate of water loss from *ahp4* mutant plants, compared with WT plants under water deficit, was partly due to reduced stomatal aperture (Figures 4d-e and S8). Similar to *ahp4* mutant, the *ahp2* single, *ahk2,3* double and *arr1,10,12* triple mutant plants also showed narrower stomatal apertures than WT (Marchadier and Hetherington, 2014; Nguyen et al., 2016). Furthermore, *ahk2,3* and *arr1,10,12* mutants exhibited higher tolerance to drought than WT (Tran et al., 2007; Kang et al., 2012; Nguyen et al., 2016). Additionally, AHP4 was reported to interact with the type-B ARR1 in the CK signaling pathway (Dortay et al., 2006), and both *AHP4* (this study) and *ARR1* (Nguyen et al., 2016; Huang et al., 2018) act as negative regulators of drought tolerance. These findings collectively suggest that loss-of-function of *AHP4* may result in weakened CK signaling, leading to reduced stomatal apertures and enhanced drought tolerance. Thus, CK signaling is an important negative regulator of stomatal activity with respect to plant adaptation to drought.

Cuticular wax is a protective barrier, containing predominantly long-chain hydrocarbons, which plays a crucial role in plant responses to various environmental stresses, including drought (Shepherd and Wynne Griffiths, 2006; Kosma et al., 2009; Seo et al., 2011; Zhu and Xiong, 2013; Lee et al., 2014). Changes in the cuticle formation, especially cuticle thickness, have been reported to be associated with drought tolerance in a range of plant species, such as *Arabidopsis thaliana*, rice and *Camelina sativa* (Aharoni et al., 2004; Shepherd and Wynne Griffiths, 2006; Seo et al., 2011; Zhu and Xiong, 2013; Lee et al., 2014; Zhu et al., 2014; Zhou et al., 2015). Adaptation of plants to drought often requires a reduction in both stomatal and non-stomatal water loss, and the latter can be achieved by increasing the thickness of the cuticles found on the surface of leaves and stems (Lee et al., 2014; Jetter and Riederer, 2016). Our results demonstrate that AHP4 negatively regulates the thickness of the cuticle layer as *ahp4* mutant plants produced thicker cuticles (Figure 5d-e); and thus, showed lower cuticle permeability than WT (Figure 5a-b). In support of this finding, transcriptional analysis also revealed that AHP4 mediates plant drought responses by down-regulating wax biosynthesis-related genes, as indicated by increased expression of the *SHN1* and *FAR6* genes in *ahp4* mutant plants under both well-watered and dehydration conditions (Figure 5f) (Aharoni et al., 2004; Cui et al., 2016). On the other hand, expression of the *DEWAX* gene in *ahp4* mutant plants was down-regulated under dehydration conditions (Figure 5f). The DEWAX belonging to the AP2/ERF-type TF family was reported to act as a negative regulator of cuticular wax biosynthesis in *Arabidopsis* as indicated by a greater wax load in the *dewax* mutant and lower wax load in the *DEWAX*-overexpressing lines as compared with that in WT (Go et al., 2014; Suh and Go, 2014). DEWAX was also shown to negatively regulate the expression of several wax biosynthesis-related genes, including *FAR6* (Go et al., 2014). As a result of enhanced wax biosynthesis, a thicker cuticle layer with reduced water permeability might help *ahp4* mutant plants maintain higher RWC (Figures 2j, 4a and S2a) than WT plants under water deficit, which is likely to be one of the major factors contributing to the greater drought tolerance of *ahp4* mutant plants.

In plants, the production of ROS can increase under drought conditions, leading to an imbalance in ROS-generation and -scavenging, which can result in oxidative damage (Mittler, 2017; Zandalinas et al., 2018; Huang et al., 2019). Therefore, a vibrant antioxidant defense is requisite to save plants from drought-induced oxidative stress (Miller et al., 2010; Zandalinas et al., 2018). A previous study reported that overproduction of endogenous CKs promoted ROS generation and decreased ROS-scavenging enzyme activities in *Arabidopsis*, resulting in enhanced salt sensitivity (Wang et al., 2015). In this study, lower ROS levels were found in *ahp4* mutant plants than WT plants under water deficit (Figure 6a-b), indicating that the mechanisms involved in ROS elimination might be activated in *ahp4* mutant. In both well-watered and water-stressed *ahp4* mutant plants, down-regulated expression of the *ferric reduction oxidase 1* (*FRO1*) gene, which encodes a ferric-chelate reductase involved in production of O_2_^•−^, was observed (Tables S2 and S3); possibly helping to maintain low O_2_^•−^ concentrations (Figure 6a) (Mittler et al., 2004). Results of enzymatic antioxidant assays confirmed that SOD, APX, GPX and GST exhibited enhanced activity levels under water deficit in *ahp4* mutant plants when compared with WT plants (Figure 6d-g). The enhanced SOD activity in the leaves of *ahp4* may help in the elimination of excessive O_2_^•−^ by enhancing the conversion of O_2_^•−^ into H_2_O_2_ (Figure 6a, d) (Liu and He, 2016). The reduced level of H_2_O_2_ in *ahp4* mutant plants corresponded to the heightened activities of APX, GST and GPX, which are involved in the removal of H_2_O_2_ produced under unfavorable stress conditions (Figure 6b, e-g). GPX and GST also play important roles in protecting the cellular membrane by removing organic peroxides and lipid hydroperoxides produced when plants undergo abiotic stresses (Miller et al., 2010; Mostofa et al., 2015). Therefore, enhanced activity of GPX and GST implies that these enzymes might significantly contribute to the protection of cellular membrane from drought-induced reactive peroxides. In addition, higher anthocyanin concentrations were found in *ahp4* mutant than WT plants under drought (Figure S10b). Since anthocyanins have been shown to function as non-enzymatic antioxidants (Nakabayashi et al., 2014; Li et al., 2017), this may also increase the capacity of *ahp4* mutant plants to limit drought-induced ROS accumulation. The increase in anthocyanin levels in *ahp4* mutant plants during drought may be explained by transcriptional regulation of the anthocyanin/flavonoid biosynthetic pathway as indicated by both the RT-qPCR and transcriptome data (Figures S4-S6, S10c). Several studies have found positive correlations between drought tolerance and anthocyanin levels in *Arabidopsis* owing to their ROS-scavenging functions (Nakabayashi et al., 2014; Nguyen et al., 2016; Li et al., 2017; Li et al., 2020a; Li et al., 2020b). The findings of the present study suggest that enhanced anthocyanin/flavonoid biosynthesis is an important drought acclimation mechanism in plants mediated by AHP4, and perhaps through ARR1, ARR10 and/or ARR12 as anthocyanin biosynthesis was reportedly enhanced in the triple *arr1,10,12* mutant under drought (Nguyen et al., 2016).

## CONCLUSIONS

Transcriptome analysis of *ahp4* mutant and WT plants has provided valuable insight into the regulatory roles of AHP4, and thus CK signaling, in plant drought adaptation (Figure 7). Loss-of-function of *APH4* altered the expression of many genes associated with plant response to water deficit, including dehydration- and/or ABA-responsive genes important for drought tolerance. In comparison with WT plants, *ahp4* mutant plants showed enhanced ABA responsiveness and photosynthetic efficiency, increased root elongation, reduced stomatal apertures, increased wax crystal accumulation and thicker cuticles. All of these factors contribute to increased water retention in *ahp4* mutant under water deficit. Additionally, under drought, *ahp4* mutant plants had lower ROS accumulation, less electrolyte leakage and lower lipid peroxidation levels. These changes indicate decreased levels of drought-induced cellular damage in *ahp4* mutant plants, which were likely due to increased expression of genes encoding enzymatic (e.g. SOD, APX, GST and GPX) and non-enzymatic (e.g. anthocyanins) antioxidants participated in antioxidant defense. The aforementioned observations collectively indicate that AHP4 has the ability to negatively regulate multiple protective mechanisms associated with drought tolerance (Figure 7). Therefore, *AHP4* is a promising candidate gene to be identified in various crops for gene editing to generate drought-tolerant crop cultivars.

## EXPERIMENTAL PROCEDURES

### Plant materials, growth and treatments

The *Arabidopsis ahp4* mutant used in this study is from the Columbia genetic background, and it was obtained from a previous work (Hutchison et al., 2006). For dehydration treatments, 21-day-old WT plants were grown on germination medium (GM) agar plates (22°C, 16-h light/8-h dark cycle, 60 µmol m^−2^ s^−1^ photon flux density) and exposed to dehydration for the indicated time periods. For ABA treatments, 21-day-old WT plants were grown on GM agar plates and treated in a solution containing 0 µM ABA (control) or 50 µM ABA for 0, 1, 2, 4 or 6 h. After the treatments, whole plant, shoot and root samples were collected in three biological replicates, and immediately frozen in liquid nitrogen and stored at - 80°C for further analyses.

### Drought tolerance assay

For examining drought-tolerant phenotypes, we followed the published gravimetric method for biomass reduction assay (Harb and Pereira, 2011; Nguyen et al., 2018), and the survival test was done in the same-tray system (Nishiyama et al., 2011). Biomass reduction of *aph4* and WT plants under drought stress relative to respective well-watered control plants was measured at 5 days after rewatering. Per genotype, 12 pots under either well-watered or drought conditions were used for measuring the plant biomass reduction. During soil-drying, the soil moisture content was recorded at five different positions per genotype using HydroSense (Campbell Scientific Australia Pty. Ltd, Australia). Photographs were taken 3 days after rewatering and removal of inflorescences from the surviving plants. For the survival test, the survival rates were calculated from three independent experiments, in which each replicate was calculated from 30 plants per genotype.

### Osmotic tolerance assay

To examine root growth under osmotic stress, 7-day-old *ahp4* mutant and WT plants were grown on GM plates, and then transferred onto 0.5 × Murashige and Skoog (MS) plates containing 1.2% agar and 0, 100, 200 or 300 mM mannitol. Primary root length of 14-day-old plants was measured after 7 days of incubation (22°C, 16-h light/ 8-h dark, 60 µmol m^−2^ s^−1^ photon flux density).

### RWC and electrolyte leakage under drought

The *ahp4* mutant and WT plants were grown and subjected to drought stress as described in the survival test in the same-tray system (Nishiyama et al., 2011). RWC and electrolyte leakage of the detached aerial portions of the plants during soil-drying were measured at 11, 12, 13 and 14 days after drought, in accordance with the methods previously described (Nishiyama et al., 2011). Five biological replicates for each genotype were used for all experimental measurements.

### ROS accumulation and MDA content

The *ahp4* mutant and WT plants were grown and subjected to drought stress as described in the survival test in the same-tray system (Nishiyama et al., 2011). ROS accumulation of *ahp4* mutant and WT plants, which were either exposed to 12 days of drought or well-watered control conditions, was determined by using nitro blue tetrazolium and diaminobenzidine staining, in accordance with the previous methods (Mostofa et al., 2015). MDA contents of *ahp4* mutant and WT plants exposed to drought for 12 and 13 days or well-watered control conditions were measured using the previous procedure (Mostofa et al., 2015). Three biological replicates of each treatment were used for measuring MDA contents.

### Anthocyanin contents and antioxidant enzyme activities

The *ahp4* mutant and WT plants were grown and subjected to drought stress as described in the survival test using the same-tray system (Nishiyama et al., 2011). After 12 and 13 days of drought stress, rosette leaves of stressed and non-stressed plants were separately collected for determining anthocyanin contents and antioxidant enzyme activities. The anthocyanin contents of freeze-dried rosette leaves of stressed and non-stressed *ahp4* mutant and WT plants were measured as previously described (Li et al., 2017). Antioxidant enzyme activities of rosette leaves of stressed and non-stressed plants were measured following previously described methods (Mostofa et al., 2015). Total soluble protein contents were measured using the Bradford method (Bradford, 1976).

### RWC, leaf temperature, stomatal density and aperture, transpiration rate and stomata conductance measurements under dehydration

To examine these characteristics, 32-day-old *ahp4* mutant and WT plants grown on soil (22°C in light period/18°C in dark period, 12-h light/12-h dark cycle, 200 µmol m^−2^ s^−1^ photon flux density, 50% relative room humidity) were exposed to dehydration for the indicated time periods. The RWC of 32-day-old *ahp4* mutant and WT plants were recorded during dehydration treatment according to the method previously described (Nishiyama et al., 2011). Five biological replicates for each genotype were used for RWC measurements. The relative leaf temperatures of 32-day-old *ahp4* mutant and WT plants during dehydration treatment were detected using the FLIR ONE camera (FLIR Systems, Wilsonville, Oregon, USA) and Thermal Analysis software. The stomatal density and dehydration-induced stomatal closure of 32-day-old *ahp4* mutant and WT plants were performed at 0 (control), 30 and 60 minutes of dehydration treatment. At the indicated time points, the stomata of the fifth leaf were embedded using Parkell Cinch hydrophilic vinyl polysiloxane (dental impression material) and Cartridge gun (Parkell Inc., Edgewood, New York, USA) for 1 h. The epidermal cells were peeled and then dried using Sally Hansen Double Duty (Sally Hansen, USA) for 1h. The epidermal cells were then used for determination of stomatal density and apertures using Leica DM750 microscope (Leica Microsystems Inc., Buffalo Grove, Illinois, USA) and ImageJ software (https://imagej.nih.gov/ij/). Five biological replicates for each genotype were used for stomatal measurements.

For transpiration rate and stomatal conductance measurements: for the dehydration samples, the whole rosette leaves of 32-day-old *ahp4* mutant and WT plants were exposed to dehydration for 30 minutes, followed by 20 minutes of dark adaptation before measurement; for the well-watered samples, plants were kept for 20 minutes under dark before measurement. The constant systems of 2-cm chamber were set during the measurement (22°C of leaf temperature, 10,000 rpm fan speed, 500 μmol s^−1^ flow rate, 400 ppm CO_2_ level). The fifth rosette leaves of 32-day-old *ahp4* mutant and WT plants were selected for determining the transpiration rate, stomata conductance at 30 minutes of dehydration or well-watered (control) conditions by using LICOR6800 system (LICOR Biosciences, Lincoln, Nebraska, USA). Three biological replicates for each genotype were used for all experimental measurements.

### The photosynthetic efficiency, chlorophyll index and NPQ under drought

The *ahp4* mutant and WT plants were grown on soil under well-watered conditions (22°C in light period/18°C in dark period, 12-h light/12-h dark cycle, 200 µmol m^−2^ s^−1^ photon flux density, 50% relative room humidity) for 35 days, and then exposed to drought stress for 14 days. The plants were kept for 20 minutes under dark before measurement. The photosynthetic efficiency (Fv/Fm), chlorophyll index and NPQ of 49-day-old *ahp4* mutant and WT plants under well-watered or drought conditions were measured using the CropReporter system (CID AgTech, Camas, Washington, USA). Four biological replicates for each genotype were used for all experimental measurements.

### Germination and root inhibition assays for evaluation of ABA responsiveness, and ABA-induced stomatal closure

Germination assay was conducted on GM medium containing 1% sucrose and various concentrations of exogenous ABA as previously described (Nishiyama et al., 2011). For root inhibition assay, 7-day-old *ahp4* mutant and WT plants grown on GM plates were transferred onto 0.5 × Murashige and Skoog (MS) plates containing 1.2% agar and 0 or 20 μM ABA. Primary root growth was measured after 4 and 7 days of incubation (22°C, 16-h light/ 8-h dark, 60 µmol m^−2^ s^−1^ photon flux density). The ABA-induced stomatal closure was performed following the previous method (Osakabe et al., 2013). Fourteen-day-old *ahp4* mutant and WT plants grown on GM plates were transferred to soil and grown for 7 additional days under well-watered conditions. Rosette leaves from 21-day-old plants were then used for determination of stomatal apertures under 0 (control), 30 and 50 μM ABA treatments (Osakabe et al., 2013).

### Chlorophyll leaching, TB staining, and determination of epicuticular wax density and cuticle thickness

For chlorophyll leaching assay, the *ahp4* mutant and WT plants were grown on soil under well-watered conditions (22°C in light period/18°C in dark period, 12-h light/12-h dark cycle, 200 µmol m^−2^ s^−1^ photon flux density, 50% relative room humidity) for 35 days. The chlorophyll leaching rates from rosette leaves of *ahp4* and WT plants were determined as previously described (Li et al., 2017). TB staining was conducted following the published procedure (Tanaka et al., 2004). The aerial portions of 35-day-old plants grown on soil were submerged into a solution containing water or 0.05% (w/v) TB for 3 h. Treated aerial portions were subsequently transferred to water and gently shaken to remove excessive TB, and were then photographed using the Leica DM750 microscope. For the assay of epicuticular wax density in stems, 14-day-old *ahp4* mutant and WT plants grown on GM plates were transferred to soil and grown for 21 additional days under well-watered conditions in the same-tray system (Nishiyama et al., 2011). The main stems of 35-day-old plants were then selected for measuring wax crystal surface by SEM (Ukitsu et al., 2007). The fifth rosette leaves of 35-day-old plants were also selected for determining cuticle layer thickness by TEM following previously reported procedures (Ukitsu et al., 2007). ImageJ software was used to measure the cuticle thickness, with eight different areas measured for each replicate.

### Gene expression analyses

Total RNA was purified using the RNeasy Plant Mini Kit (Qiagen, Hilden, Germany). The cDNA synthesis and RT-qPCR were conducted according to previous methods (Le et al., 2011). *UBQ10* was used as a reference gene for RT-qPCR data analysis. Gene-specific primers used for RT-qPCR are presented in Table S6.

### Dehydration sampling and microarray analysis

Fourteen-day-old *ahp4* mutant and WT plants grown on GM plates were transferred to soil and grown for 10 additional days under well-watered conditions. The aerial portions of 24-day-old plants were then subjected to dehydration treatments as previously described (Ha et al., 2014). Rosette leaves of *ahp4* mutant and WT plants treated by dehydration for 0, 2 and 5 h were collected in three biological repeats, and were then used for transcriptome analysis using the *Arabidopsis* Oligo 44K DNA microarray (Version 4.0, Agilent Technology) (Nishiyama et al., 2012). To search for DEGs, the criteria of |fold-change| ≥ 2 and a false discovery rate corrected *P*-value (q-value) of < 0.05 were used. The detailed protocol and raw microarray data have been deposited in the Gene Expression Omnibus database (GSE95614). ClustVis (https://biit.cs.ut.ee/clustvis/) and MapMan (http://mapman.gabipd.org) were used to analyze the data.

### Statistical analysis

Analysis of variance (ANOVA) or Student’s *t*-test was used for data analysis. Different superscripted letters within the column reveal statistically significant differences between the two genotypes and among the treatments as determined by Duncan’s multiple range test (using IBM SPSS software package 21.0). Asterisks demonstrate significant differences as assessed by the Student’s *t*-test (\**P* < 0.05, \*\**P* < 0.01, \*\*\**P* < 0.001).

## ACCESSION NUMBERS

The data reported in this paper have been deposited in the Gene Expression Omnibus (GEO) database, www.ncbi.nlm.nih.gov/geo (accession no. GSE95614). ID: chienhavan. Password: Hana18289.

## Supporting information

Results of the comparative microarray analysis of leaves of ahp4 mutant and wild-type plants under well-watered and dehydration conditions.

List of up- and down-regulated genes in M-C/W-C comparison

List of up- and down-regulated genes in various comparisons

Venn analysis of differentially expressed gene sets derived from various comparisons

List of photosynthesis-related genes in ahp4 and wild-type (WT) leaves under normal and dehydration conditions

Primers used in RT-qPCR

## ACKNOWLEDGMENTS

The authors would like to thank Dr. Sona Pandey (Donald Danforth Plant Science Center), and Rie Nishiyama and Yu Li (RIKEN CSRS) for their support in this research, as well as Mayumi Wakazaki (RIKEN CSRS) for supporting the TEM observations, and Audrey Dodds (Donald Danforth Plant Science Center) for editing the English.

## AUTHOR CONTRIBUTIONS

L.-S.P.T. designed the research; C.V.H., K.H.N., M.G.M., C.D.T., Y.W., W.L., Y.O., M.S., K.T., M.T., C.A., R.Z. and M.S. performed the research; K.T., and M.S. contributed research materials, reagents and analytic tools; C.V.H., M.G.M., C.A and R.Z. analyzed the data with the input of D.J.B. and L.-S.P.T.; and C.V.H., D.J.B. and L.-S.P.T. critically interpreted the study and wrote the manuscript.

## SUPPORTING INFORMATION

**Figure S1.** A representative assay of root growth of *ahp4* mutant and wild-type (WT) plants under normal and mannitol-induced osmotic stress conditions.

**Figure S2.** Comparative transcriptome analysis of *ahp4* mutant and wild-type (WT) plants exposed to a dehydration stress.

**Figure S3.** Confirmation of microarray data by real-time quantitative PCR (RT-qPCR) analysis.

**Figure S4.** MapMan-based analysis of differentially expressed genes identified in *ahp4* mutant versus wild-type (WT) plants under normal (M-C/W-C comparison) and dehydration (M-D/W-D comparison) conditions.

**Figure S5.** Metabolism-related overview of differentially expressed genes derived from *ahp4* versus wild-type under normal and dehydration conditions using MapMan.

**Figure S6.** Secondary metabolism-related overview of differentially expressed genes identified in *ahp4* versus wild-type under normal and dehydration conditions using MapMan.

**Figure S7.** Stomatal conductance and leaf transpiration rates of *ahp4* mutant and wild-type (WT) plants under well-watered and dehydration conditions.

**Figure S8.** Comparison of stomatal apertures of *ahp4* mutant and wild-type (WT) plants under abscisic acid (ABA) treatment.

**Figure S9.** Germination assay of *ahp4* mutant and wild-type (WT) plants on medium supplemented with different concentrations of abscisic acid (ABA), and *AHP4* expression in WT plants treated with ABA.

**Figure S10.** Anthocyanin contents and expression of anthocyanin/flavonoid-related genes in *ahp4* mutant and wild-type (WT) plants under drought.

**Table S1.** Results of the comparative microarray analysis of leaves of *ahp4* mutant and wild-type plants under well-watered and dehydration conditions.

**Table S2.** List of up- and down-regulated genes in M-C/W-C comparison (|fold-change| ≥ 2; q-value < 0.05).

**Table S3.** List of up- and down-regulated genes in various comparisons (|fold-change| ≥ 2; q-value < 0.05).

**Table S4.** Venn analysis of differentially expressed gene sets derived from various comparisons.

**Table S5.** List of photosynthesis-related genes in *ahp4* and wild-type (WT) leaves under normal and dehydration conditions.

**Table S6.** Primers used in RT-qPCR.

## REFERENCES

Abdeen A, Schnell J, Miki B (2010) Transcriptome analysis reveals absence of unintended effects in drought-tolerant transgenic plants overexpressing the transcription factor ABF3. BMC Genomics 11: 69.

Abdelrahman M, Burritt DJ, Tran LP (2018) The use of metabolomic quantitative trait locus mapping and osmotic adjustment traits for the improvement of crop yields under environmental stresses. Semin Cell Dev Biol 83: 86–94.

Abdelrahman M, El-Sayed M, Jogaiah S, Burritt DJ, Tran LP (2017) The “STAY-GREEN” trait and phytohormone signaling networks in plants under heat stress. Plant Cell Rep 36: 1009–1025.

Aharoni A, Dixit S, Jetter R, Thoenes E, van Arkel G, Pereira A (2004) The SHINE clade of AP2 domain transcription factors activates wax biosynthesis, alters cuticle properties, and confers drought tolerance when overexpressed in Arabidopsis. Plant Cell 16: 2463–2480.

Bedi S, Chaudhuri RN (2018) Transcription factor ABI3 auto-activates its own expression during dehydration stress response. FEBS Lett 592: 2594–2611.

Bedi S, Sengupta S, Ray A, Nag Chaudhuri R (2016) ABI3 mediates dehydration stress recovery response in Arabidopsis thaliana by regulating expression of downstream genes. Plant Sci 250: 125–140.

Bradford MM (1976) A rapid and sensitive method for the quantitation of microgram quantities of protein utilizing the principle of protein-dye binding. Anal Biochem 72: 248–254.

Choudhury FK, Rivero RM, Blumwald E, Mittler R (2017) Reactive oxygen species, abiotic stress and stress combination. Plant J 90: 856–867.

Cortleven A, Leuendorf JE, Frank M, Pezzetta D, Bolt S, Schmulling T (2018) Cytokinin action in response to abiotic and biotic stress in plants. Plant Cell Environ 42: 998–1018.

Cui F, Brosche M, Lehtonen MT, Amiryousefi A, Xu E, Punkkinen M, Valkonen JP, Fujii H, Overmyer K (2016) Dissecting abscisic acid signaling pathways involved in cuticle formation. Mol Plant 9: 926–938.

Daryanto S, Wang L, Jacinthe PA (2016) Global synthesis of drought effects on maize and wheat production. PLoS One 11: e0156362.

Ding S, Zhang B, Qin F (2015) Arabidopsis RZFP34/CHYR1, a Ubiquitin E3 ligase, regulates stomatal movement and drought tolerance via SnRK2.6-mediated phosphorylation. Plant Cell 27: 3228–3244.

Dong T, Xu ZY, Park Y, Kim DH, Lee Y, Hwang I (2014) Abscisic acid uridine diphosphate glucosyltransferases play a crucial role in abscisic acid homeostasis in Arabidopsis. Plant Physiol 165: 277–289.

Dortay H, Mehnert N, Burkle L, Schmulling T, Heyl A (2006) Analysis of protein interactions within the cytokinin-signaling pathway of Arabidopsis thaliana. FEBS J 273: 4631–4644.

Vanjildorj E, Bae TW, Riu KZ, Kim SY, Lee HY (2005) Overexpression of Arabidopsis ABF3 gene enhances tolerance to drought and cold in transgenic lettuce (Lactuca sativa). Plant Cell Tiss Org 83.: 41–50.

Farooq MA, Niazi AK, Akhtar J, Saifullah, Farooq M, Souri Z, Karimi N, Rengel Z (2019) Acquiring control: The evolution of ROS-induced oxidative stress and redox signaling pathways in plant stress responses. Plant Physiol Biochem 141: 353–369.

Feng W, Lindner H, Robbins NE 2nd, Dinneny JR (2016) Growing out of stress: The role of cell- and organ-scale growth control in plant water-stress responses. Plant Cell 28: 1769–1782.

Fujita M, Fujita Y, Maruyama K, Seki M, Hiratsu K, Ohme-Takagi M, Tran LS, Yamaguchi-Shinozaki K, Shinozaki K (2004) A dehydration-induced NAC protein, RD26, is involved in a novel ABA-dependent stress-signaling pathway. Plant J 39: 863–876.

Fujita Y, Fujita M, Satoh R, Maruyama K, Parvez MM, Seki M, Hiratsu K, Ohme-Takagi M, Shinozaki K, Yamaguchi-Shinozaki K (2005) AREB1 is a transcription activator of novel ABRE-dependent ABA signaling that enhances drought stress tolerance in Arabidopsis. Plant Cell 17: 3470–3488.

Fujita Y, Yoshida T, Yamaguchi-Shinozaki K (2013) Pivotal role of the AREB/ABF-SnRK2 pathway in ABRE-mediated transcription in response to osmotic stress in plants. Physiol Plant 147: 15–27.

Go YS, Kim H, Kim HJ, Suh MC (2014) Arabidopsis cuticular wax biosynthesis is negatively regulated by the DEWAX gene encoding an AP2/ERF-type transcription factor. Plant Cell 26: 1666–1680.

Ha CV, Leyva-Gonzalez MA, Osakabe Y, Tran UT, Nishiyama R, Watanabe Y, Tanaka M, Seki M, Yamaguchi S, Dong NV, Yamaguchi-Shinozaki K, Shinozaki K, Herrera-Estrella L, Tran LS (2014) Positive regulatory role of strigolactone in plant responses to drought and salt stress. Proc Natl Acad Sci U S A 111: 851–856.

Ha S, Vankova R, Yamaguchi-Shinozaki K, Shinozaki K, Tran LS (2012) Cytokinins: metabolism and function in plant adaptation to environmental stresses. Trends Plant Sci 17: 172–179.

Han YJ, Cho KC, Hwang OJ, Choi YS, Shin AY, Hwang I, Kim JI (2012) Overexpression of an Arabidopsis beta-glucosidase gene enhances drought resistance with dwarf phenotype in creeping bentgrass. Plant Cell Rep 31: 1677–1686.

Harb A, Pereira A (2011) Screening Arabidopsis genotypes for drought stress resistance. Methods Mol Biol 678: 191–198

Huang H, Ullah F, Zhou DX, Yi M, Zhao Y (2019) Mechanisms of ROS regulation of plant development and stress responses. Front Plant Sci 10: 800.

Huang X, Hou L, Meng J, You H, Li Z, Gong Z, Yang S, Shi Y (2018) The antagonistic action of abscisic acid and cytokinin signaling mediates drought stress response in Arabidopsis. Mol Plant 11: 970–982.

Hutchison CE, Li J, Argueso C, Gonzalez M, Lee E, Lewis MW, Maxwell BB, Perdue TD, Schaller GE, Alonso JM, Ecker JR, Kieber JJ (2006) The Arabidopsis histidine phosphotransfer proteins are redundant positive regulators of cytokinin signaling. Plant Cell 18: 3073–3087.

Jetter R, Riederer M (2016) Localization of the transpiration barrier in the epi- and intracuticular waxes of eight plant species: Water transport resistances are associated with fatty acyl rather than alicyclic components. Plant Physiol 170: 921–934.

Kang JY, Choi HI, Im MY, Kim SY (2002) Arabidopsis basic leucine zipper proteins that mediate stress-responsive abscisic acid signaling. Plant Cell 14: 343–357.

Kang NY, Cho C, Kim NY, Kim J (2012) Cytokinin receptor-dependent and receptor-independent pathways in the dehydration response of Arabidopsis thaliana. J Plant Physiol 169: 1382–1391.

Kerr TC, Abdel-Mageed H, Aleman L, Lee J, Payton P, Cryer D, Allen RD (2017) Ectopic expression of two AREB/ABF orthologs increase dehydration tolerance in cotton (Gossypium hirsutum). Plant Cell Environ 41: 898–907.

Keshishian EA, Rashotte AM (2015) Plant cytokinin signalling. Essays Biochem 58: 13–27.

Kieber JJ, Schaller GE (2018) Cytokinin signaling in plant development. Development 145: dev149344.

Kim HJ, Kieber JJ, Schaller GE (2013) The rice F-box protein KISS ME DEADLY2 functions as a negative regulator of cytokinin signalling. Plant Signal Behav 8: e26434.

Kim JB, Kang JY, Kim SY (2004) Over-expression of a transcription factor regulating ABA-responsive gene expression confers multiple stress tolerance. Plant Biotechnol J 2: 459–466.

Kosma DK, Bourdenx B, Bernard A, Parsons EP, Lu S, Joubes J, Jenks MA (2009) The impact of water deficiency on leaf cuticle lipids of Arabidopsis. Plant Physiol 151: 1918–1929.

Kumar MN, Verslues PE (2015) Stress physiology functions of the Arabidopsis histidine kinase cytokinin receptors. Physiol Plant 154: 369–380.

Kuromori T, Seo M, Shinozaki K (2018) ABA transport and plant water stress responses. Trends Plant Sci 23: 513–522.

Lamaoui M, Jemo M, Datla R, Bekkaoui F (2018) Heat and drought stresses in crops and approaches for their mitigation. Front Chem 6: 26.

Le DT, Nishiyama R, Watanabe Y, Mochida K, Yamaguchi-Shinozaki K, Shinozaki K, Tran LS (2011) Genome-wide expression profiling of soybean two-component system genes in soybean root and shoot tissues under dehydration stress. DNA Res 18: 17–29.

Lee KH, Piao HL, Kim HY, Choi SM, Jiang F, Hartung W, Hwang I, Kwak JM, Lee IJ (2006) Activation of glucosidase via stress-induced polymerization rapidly increases active pools of abscisic acid. Cell 126: 1109–1120.

Lee SB, Kim H, Kim RJ, Suh MC (2014) Overexpression of Arabidopsis MYB96 confers drought resistance in Camelina sativa via cuticular wax accumulation. Plant Cell Rep 33: 1535–1546.

Li P, Li YJ, Zhang FJ, Zhang GZ, Jiang XY, Yu HM, Hou BK (2017) The Arabidopsis UDP-glycosyltransferases UGT79B2 and UGT79B3, contribute to cold, salt and drought stress tolerance via modulating anthocyanin accumulation. Plant J 89: 85–103.

Li W, Herrera-Estrella L, Tran LP (2019) Do cytokinins and strigolactones crosstalk during drought adaptation? Trends Plant Sci 24: 669–672.

Li W, Herrera-Estrella L, Tran LS (2016) The yin-yang of cytokinin homeostasis and drought acclimation/adaptation. Trends Plant Sci 21: 548–550

Li W, Nguyen KH, Chu HD, Ha CV, Watanabe Y, Osakabe Y, Leyva-Gonzalez MA, Sato M, Toyooka K, Voges L, Tanaka M, Mostofa MG, Seki M, Seo M, Yamaguchi S, Nelson DC, Tian C, Herrera-Estrella L, Tran LP (2017) The karrikin receptor KAI2 promotes drought resistance in Arabidopsis thaliana. PLoS Genet 13: e1007076.

Li W, Nguyen KH, Chu HD, Watanabe Y, Osakabe Y, Sato M, Toyooka K, Seo M, Tian L, Tian C, Yamaguchi S, Tanaka M, Seki M, Tran LP (2020). Comparative functional analyses of DWARF14 and KARRIKIN INSENSITIVE 2 in drought adaptation of Arabidopsis thaliana. Plant J 03:111–127.

Li W, Nguyen KH, Ha CV, Watanabe Y, Tran LP (2019) Crosstalk between the cytokinin and MAX2 signaling pathways in growth and callus formation of Arabidopsis thaliana. Biochem Biophys Res Commun 511: 300–306.

Li W, Nguyen KH, Tran CD, Watanabe Y, Tian C, Yin X, Li K, Yang Y, Guo J, Miao Y, Yamaguchi S, Tran LP (2020). Negative roles of strigolactone-related SMXL6, 7 and 8 proteins in drought resistance in Arabidopsis. Biomolecules 10:607.

Liu Y, He C (2016) Regulation of plant reactive oxygen species (ROS) in stress responses: learning from AtRBOHD. Plant Cell Rep 35: 995–1007.

Lotkowska ME, Tohge T, Fernie AR, Xue GP, Balazadeh S, Mueller-Roeber B (2015) The Arabidopsis transcription factor MYB112 promotes anthocyanin formation during salinity and under high light stress. Plant Physiol 169: 1862–1880.

Marchadier E, Hetherington AM (2014) Involvement of two-component signalling systems in the regulation of stomatal aperture by light in Arabidopsis thaliana. New Phytol 203: 462–468.

Miller G, Suzuki N, Ciftci-Yilmaz S, Mittler R (2010) Reactive oxygen species homeostasis and signalling during drought and salinity stresses. Plant Cell Environ 33: 453–467

Mittler R (2017) ROS are good. Trends Plant Sci 22: 11–19. doi: 10.1111/j.1365-3040.2009.02041.x.

Mittler R, Vanderauwera S, Gollery M, Van Breusegem F (2004) Reactive oxygen gene network of plants. Trends Plant Sci 9: 490–498.

Mostofa MG, Hossain MA, Fujita M, Tran LS (2015) Physiological and biochemical mechanisms associated with trehalose-induced copper-stress tolerance in rice. Sci Rep 5: 11433.

Mostofa MG, Li W, Nguyen KH, Fujita M, Tran LP (2018) Strigolactones in plant adaptation to abiotic stresses: An emerging avenue of plant research. Plant Cell Environ 41: 2227–2243.

Nakabayashi R, Yonekura-Sakakibara K, Urano K, Suzuki M, Yamada Y, Nishizawa T, Matsuda F, Kojima M, Sakakibara H, Shinozaki K, Michael AJ, Tohge T, Yamazaki M, Saito K (2014) Enhancement of oxidative and drought tolerance in Arabidopsis by overaccumulation of antioxidant flavonoids. Plant J 77: 367–379.

Nguyen KH, Ha CV, Nishiyama R, Watanabe Y, Leyva-Gonzalez MA, Fujita Y, Tran UT, Li W, Tanaka M, Seki M, Schaller GE, Herrera-Estrella L, Tran LS (2016) Arabidopsis type B cytokinin response regulators ARR1, ARR10, and ARR12 negatively regulate plant responses to drought. Proc Natl Acad Sci U S A 113: 3090–3095.

Nguyen KH, Mostofa MG, Li W, Ha CV, Watanabe Y, Le DT, Nguyen TP, Tran LS (2018) The soybean transcription factor GmNAC085 enhances drought tolerance in Arabidopsis. Environ Exp Bot 151: 12–20.

Nguyen NH, Kim JH, Kwon J, Jeong CY, Lee W, Lee D, Hong SW, Lee H (2016) Characterization of Arabidopsis thaliana FLAVONOL SYNTHASE 1 (FLS1) - overexpression plants in response to abiotic stress. Plant Physiol Biochem 103: 133–142.

Nilson SE, Assmann SM (2007) The control of transpiration. Insights from Arabidopsis. Plant Physiol 143: 19–27.

Nishiyama R, Le DT, Watanabe Y, Matsui A, Tanaka M, Seki M, Yamaguchi-Shinozaki K, Shinozaki K, Tran LS (2012) Transcriptome analyses of a salt-tolerant cytokinin-deficient mutant reveal differential regulation of salt stress response by cytokinin deficiency. PLoS One 7: e32124.

Nishiyama R, Watanabe Y, Fujita Y, Le DT, Kojima M, Werner T, Vankova R, Yamaguchi-Shinozaki K, Shinozaki K, Kakimoto T, Sakakibara H, Schmulling T, Tran LS (2011) Analysis of cytokinin mutants and regulation of cytokinin metabolic genes reveals important regulatory roles of cytokinins in drought, salt and abscisic acid responses, and abscisic acid biosynthesis. Plant Cell 23: 2169–2183.

Nishiyama R, Watanabe Y, Leyva-Gonzalez MA, Ha CV, Fujita Y, Tanaka M, Seki M, Yamaguchi-Shinozaki K, Shinozaki K, Herrera-Estrella L, Tran LS (2013) Arabidopsis AHP2, AHP3, and AHP5 histidine phosphotransfer proteins function as redundant negative regulators of drought stress response. Proc Natl Acad Sci U S A 110: 4840–4845.

Oh SJ, Song SI, Kim YS, Jang HJ, Kim SY, Kim M, Kim YK, Nahm BH, Kim JK (2005) Arabidopsis CBF3/DREB1A and ABF3 in transgenic rice increased tolerance to abiotic stress without stunting growth. Plant Physiol 138: 341–351.

Osakabe Y, Arinaga N, Umezawa T, Katsura S, Nagamachi K, Tanaka H, Ohiraki H, Yamada K, Seo SU, Abo M, Yoshimura E, Shinozaki K, Yamaguchi-Shinozaki K (2013) Osmotic stress responses and plant growth controlled by potassium transporters in Arabidopsis. Plant Cell 25: 609–624.

Osakabe Y, Osakabe K, Shinozaki K, Tran LS (2014) Response of plants to water stress. Front Plant Sci 5: 86.

Osakabe Y, Yamaguchi-Shinozaki K, Shinozaki K, Tran LS (2014) ABA control of plant macroelement membrane transport systems in response to water deficit and high salinity. New Phytol 202: 35–49.

Pekarova B, Szmitkowska A, Dopitova R, Degtjarik O, Zidek L, Hejatko J (2016) Structural aspects of multistep phosphorelay-mediated signaling in plants. Mol Plant 9: 71–85.

Peleg Z, Blumwald E (2011) Hormone balance and abiotic stress tolerance in crop plants. Curr Opin Plant Biol 14: 290–295.

Prerostova S, Dobrev PI, Gaudinova A, Knirsch V, Korber N, Pieruschka R, Fiorani F, Brzobohaty B, Cerny M, Spichal L, Humplik J, Vanek T, Schurr U, Vankova R (2018) Cytokinins: Their impact on molecular and growth responses to drought stress and recovery in Arabidopsis. Front Plant Sci 9: 655.

Pruthvi V, Narasimhan R, Nataraja KN (2014) Simultaneous expression of abiotic stress responsive transcription factors, AtDREB2A, AtHB7 and AtABF3 improves salinity and drought tolerance in peanut (Arachis hypogaea L.). PLoS One 9: e111152.

Ramireddy E, Hosseini SA, Eggert K, Gillandt S, Gnad H, von Wiren N, Schmulling T(2018) Root engineering in barley: Increasing cytokinin degradation produces a larger root system, mineral enrichment in the shoot and improved drought tolerance. Plant Physiol 177: 1078–1095.

Riefler M, Novak O, Strnad M, Schmulling T (2006) Arabidopsis cytokinin receptor mutants reveal functions in shoot growth, leaf senescence, seed size, germination, root development, and cytokinin metabolism. Plant Cell 18: 40–54.

Rohde A, Kurup S, Holdsworth M (2000). ABI3 emerges from the seed. Trends Plant Sci 5: 418–419.

Romanov GA, Lomin SN, Schmulling T (2018) Cytokinin signaling: from the ER or from the PM? That is the question! New Phytol 218: 41–53.

Roscoe TT, Guilleminot J, Bessoule JJ, Berger F, Devic M (2015). Complementation of seed maturation phenotypes by ectopic expression of ABSCISIC ACID INSENSITIVE3, FUSCA3 and LEAFY COTYLEDON2 in Arabidopsis. Plant Cell Physiol 56:1215–28.

Sah SK, Reddy KR, Li J (2016) Abscisic acid and abiotic stress tolerance in crop plants. Front Plant Sci 7: 571.

Saradadevi R, Palta JA, Siddique KHM (2017) ABA-mediated stomatal response in regulating water use during the development of terminal drought in wheat. Front Plant Sci 8: 1251.

Seo PJ, Lee SB, Suh MC, Park MJ, Go YS, Park CM (2011) The MYB96 transcription factor regulates cuticular wax biosynthesis under drought conditions in Arabidopsis. Plant Cell 23: 1138–1152.

Shepherd T, Wynne Griffiths D (2006) The effects of stress on plant cuticular waxes. New Phytol 171: 469–499.

Sieber P, Schorderet M, Ryser U, Buchala A, Kolattukudy P, Metraux JP, Nawrath C (2000) Transgenic Arabidopsis plants expressing a fungal cutinase show alterations in the structure and properties of the cuticle and postgenital organ fusions. Plant Cell 12: 721–738.

Suh MC, Go YS (2014) DEWAX-mediated transcriptional repression of cuticular wax biosynthesis in Arabidopsis thaliana. Plant Signal Behav 9: e29463.

Tanaka T, Tanaka H, Machida C, Watanabe M, Machida Y (2004) A new method for rapid visualization of defects in leaf cuticle reveals five intrinsic patterns of surface defects in Arabidopsis. Plant J 37: 139–146.

Tran LS, Urao T, Qin F, Maruyama K, Kakimoto T, Shinozaki K, Yamaguchi-Shinozaki K (2007) Functional analysis of AHK1/ATHK1 and cytokinin receptor histidine kinases in response to abscisic acid, drought, and salt stress in Arabidopsis. Proc Natl Acad Sci U S A 104: 20623–20628.

Verma V, Ravindran P, Kumar PP (2016) Plant hormone-mediated regulation of stress responses. BMC Plant Biol 16: 86.

Verslues PE, Agarwal M, Katiyar-Agarwal S, Zhu J, Zhu JK (2006) Methods and concepts in quantifying resistance to drought, salt and freezing, abiotic stresses that affect plant water status. Plant J 45: 523–539.

Wang Y, Li L, Ye T, Zhao S, Liu Z, Feng YQ, Wu Y (2011) Cytokinin antagonizes ABA suppression to seed germination of Arabidopsis by downregulating ABI5 expression. Plant J 68: 249–261.

Wang Y, Shen W, Chan Z, Wu Y (2015) Endogenous cytokinin overproduction modulates ROS homeostasis and decreases salt stress resistance in Arabidopsis Thaliana. Front Plant Sci 6: 1004.

Wang Z, Li G, Sun H, Ma L, Guo Y, Zhao Z, Gao H, Mei L (2018) Effects of drought stress on photosynthesis and photosynthetic electron transport chain in young apple tree leaves. Biol Open 7: pii: bio035279. doi: 10.1242/bio.035279.

Wang Z, Su G, Li M, Ke Q, Kim SY, Li H, Huang J, Xu B, Deng XP, Kwak SS (2016) Overexpressing Arabidopsis ABF3 increases tolerance to multiple abiotic stresses and reduces leaf size in alfalfa. Plant Physiol Biochem 109: 199–208.

Werner T, Nehnevajova E, Kollmer I, Novak O, Strnad M, Kramer U, Schmulling T (2010) Root-specific reduction of cytokinin causes enhanced root growth, drought tolerance, and leaf mineral enrichment in Arabidopsis and tobacco. Plant Cell 22: 3905–3920.

Wybouw B, De Rybel B (2019) Cytokinin - a developing story. Trends Plant Sci 24: 177–185.

Xie X, He Z, Chen N, Tang Z, Wang Q, Cai Y (2019) The roles of environmental factors in regulation of oxidative stress in plant. Biomed Res Int 2019: 9732325.

Yeats TH, Rose JK (2013) The formation and function of plant cuticles. Plant Physiol 163: 5–20.

Yoshida T, Fujita Y, Maruyama K, Mogami J, Todaka D, Shinozaki K, Yamaguchi-Shinozaki K (2015) Four Arabidopsis AREB/ABF transcription factors function predominantly in gene expression downstream of SnRK2 kinases in abscisic acid signalling in response to osmotic stress. Plant Cell Environ 38: 35–49.

Zandalinas SI, Mittler R, Balfagon D, Arbona V, Gomez-Cadenas A (2018) Plant adaptations to the combination of drought and high temperatures. Physiol Plant 162: 2–12.

Zandkarimi H, Ebadi A, Salami SA, Alizade H, Baisakh N (2015) Analyzing the expression profile of AREB/ABF and DREB/CBF genes under drought and salinity stresses in grape (Vitis vinifera L.). PLoS One 10: e0134288.

Zhou X, Li L, Xiang J, Gao G, Xu F, Liu A, Zhang X, Peng Y, Chen X, Wan X (2015) OsGL1-3 is involved in cuticular wax biosynthesis and tolerance to water deficit in rice. PLoS One 10: e116676.

Zhu L, Guo J, Zhu J, Zhou C (2014) Enhanced expression of EsWAX1 improves drought tolerance with increased accumulation of cuticular wax and ascorbic acid in transgenic Arabidopsis. Plant Physiol Biochem 75: 24–35.

Zhu X, Xiong L (2013) Putative megaenzyme DWA1 plays essential roles in drought resistance by regulating stress-induced wax deposition in rice. Proc Natl Acad Sci U S A 110: 17790–17795.

Zwack PJ, Rashotte AM (2015) Interactions between cytokinin signalling and abiotic stress responses. J Exp Bot 66: 4863–4871.

