## Supplementary material for "The histidine phosphotransfer AHP4 plays a negative role in *Arabidopsis* plant response to drought"

**SUPPLEMENTAL DATA**

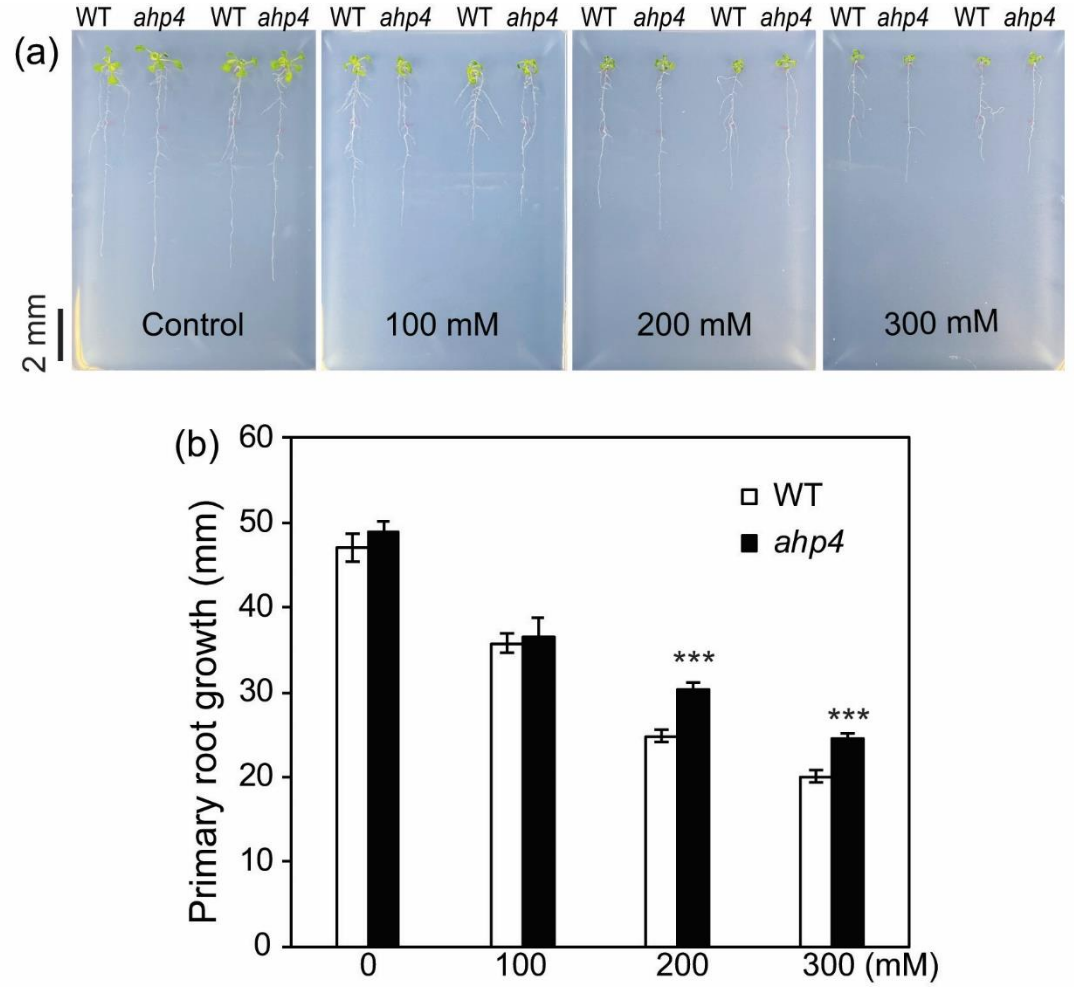


**Figure S1.** A representative assay of root growth of *ahp4* mutant and wild-type (WT) plants under normal and mannitol-induced osmotic stress conditions. (a) Seven-day-old *ahp4* and WT plants grown on germination medium were transferred onto 0.5 × MS plates (1.2% w/v agar) with or without mannitol and grown for 7 additional days. (b) Primary root growth of 7-day-old *ahp4* and WT plants grown under normal and mannitol conditions for 7 additional days. Data present the means and standard errors ($n$ = 18 plants/genotype). Asterisks indicate significant differences between the two genotypes as determined by a Student's $t$-test (***$P < 0.001$).

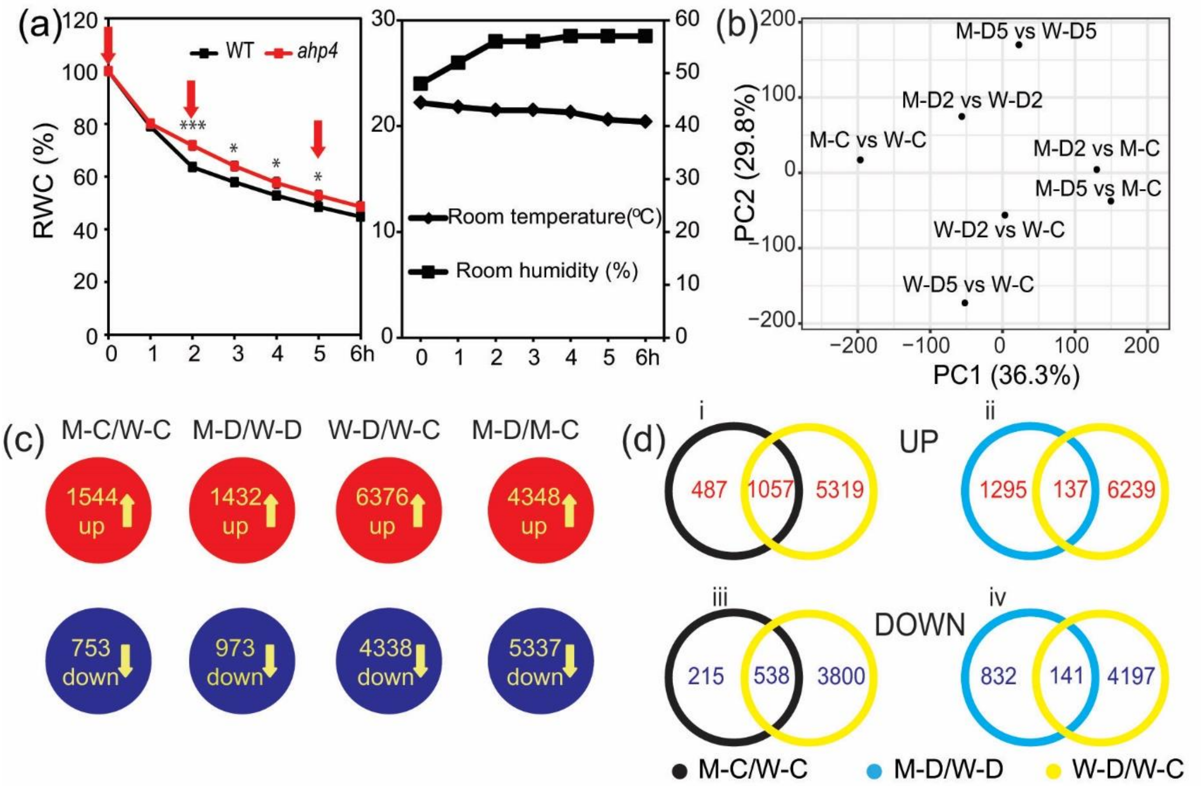


**Figure S2.** Comparative transcriptome analysis of *ahp4* mutant and wild-type (WT) plants exposed to a dehydration stress. (a) Fourteen-day-old *ahp4* and WT plants grown on GM plates were transferred to soil and grown for 10 additional days under well-watered conditions. The aerial portions of 24-day-old *ahp4* and WT plants were subjected to a dehydration stress and recorded the relative water content (RWC). Data represent the means and standard errors ($n$ = 5/genotype). Asterisks indicate significant differences between the two genotypes as determined by a Student's $t$-test (*$P < 0.05$, ***$P < 0.001$). Rosette leaves, which were collected at 0, 2, and 5 hours (h) (arrows) of dehydration treatment, were used for microarray analysis. Room temperature and relative room humidity were recorded during the dehydration treatment. (b) Principal component (PC) analysis plots of *ahp4* and WT in various comparisons. (c) Diagrams showing the compilation of genes with changed expression in each comparison. Data were extracted from the results of three independent microarray experiments. (d) Venn diagram analysis showing the overlapping and non-overlapping up- and down-regulated gene sets. M-C/W-C, *ahp4* well-watered control 0 h versus WT well-watered control 0 h; M-D2/W-D2, *ahp4* dehydrated 2 h versus WT dehydrated 2 h; M-D5/W-D5, *ahp4* dehydrated 5 h versus WT dehydrated 5 h; W-D2/W-C, WT dehydrated 2 h versus WT well-watered control 0 h; W-D5/W-C, WT dehydrated 5 h versus WT well-watered control 0 h; M-D2/M-C, *ahp4* dehydrated 2 h versus *ahp4* well-watered control 0 h; M-D5/M-C, *ahp4* dehydrated 5 h versus *ahp4* well-watered control 0 h; M-D/W-D, *ahp4* dehydrated 2 and/or 5 h versus WT dehydrated 2 and/or 5 h; M-D/M-C, *ahp4* dehydrated 2 and/or 5 h versus *ahp4* well-watered control 0 h; W-D/W-C, WT dehydrated 2 and/or 5 h versus WT well-watered control 0 h.

(a)

| AGI | Gene description | M-C / W-C | | M-D2 / W-D2 | | M-D5 / W-D5 | |
|---|---|---|---|---|---|---|---|
| | | q-value | Fold change | q-value | Fold change | q-value | Fold change |
| *AT4G34000* | *ABF3* | 0.5779 | 0.6828 | 0.4793 | 3.4051 | 0.0202 | 5.1985 |
| *AT3G21600* | Dehydration-associated protein-like protein | 0.0109 | 2.1202 | 0.0878 | 1.3197 | 0.1450 | 0.6364 |
| *AT4G37410* | *CYP81F4* | 0.0285 | 9.8801 | 0.3303 | 0.5558 | 0.4122 | 0.6257 |
| *AT2G40610* | *EXPA8* | 0.0385 | 0.4844 | 0.5403 | 1.2889 | 0.7679 | 0.7244 |
| *AT5G66390* | *P72* | 0.0048 | 4.7093 | 0.6973 | 1.2666 | 0.3435 | 1.3136 |

(b)

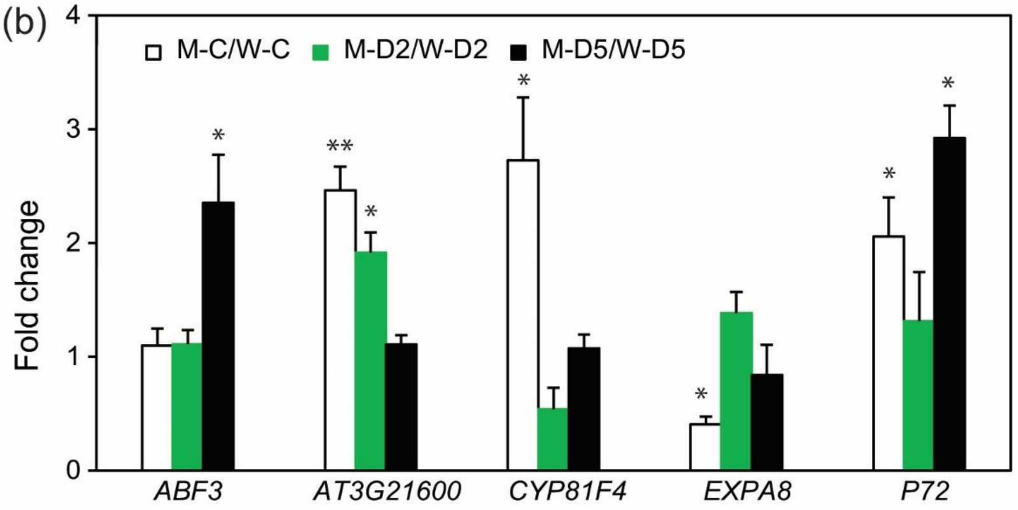


**Figure S3.** Confirmation of microarray data by real-time quantitative PCR (RT-qPCR) analysis. Five genes were selected, and their expression patterns were examined by RT-qPCR in the leaf samples of non-dehydrated and dehydrated *ahp4* mutant and wild-type (WT) plants to validate the microarray data. (a) Fold changes were extracted from microarray analysis. Red and blue colors indicate up- and down-regulated genes, respectively. (b) Fold changes were obtained from RT-qPCR analysis of three independent biological replicates. Bars represent means and standard errors ($n = 3$). Asterisks indicate significant differences as determined by a Student's *t*-test (*$P < 0.05$, **$P < 0.01$). M-C/W-C, *ahp4*-well-watered control-0 h versus WT-well-watered control-0 h; M-D2/W-D2, *ahp4*-dehydrated-2 h versus WT-dehydrated-2 h; M-D5/W-D5, *ahp4*-dehydrated-5 h versus WT-dehydrated-5 h.

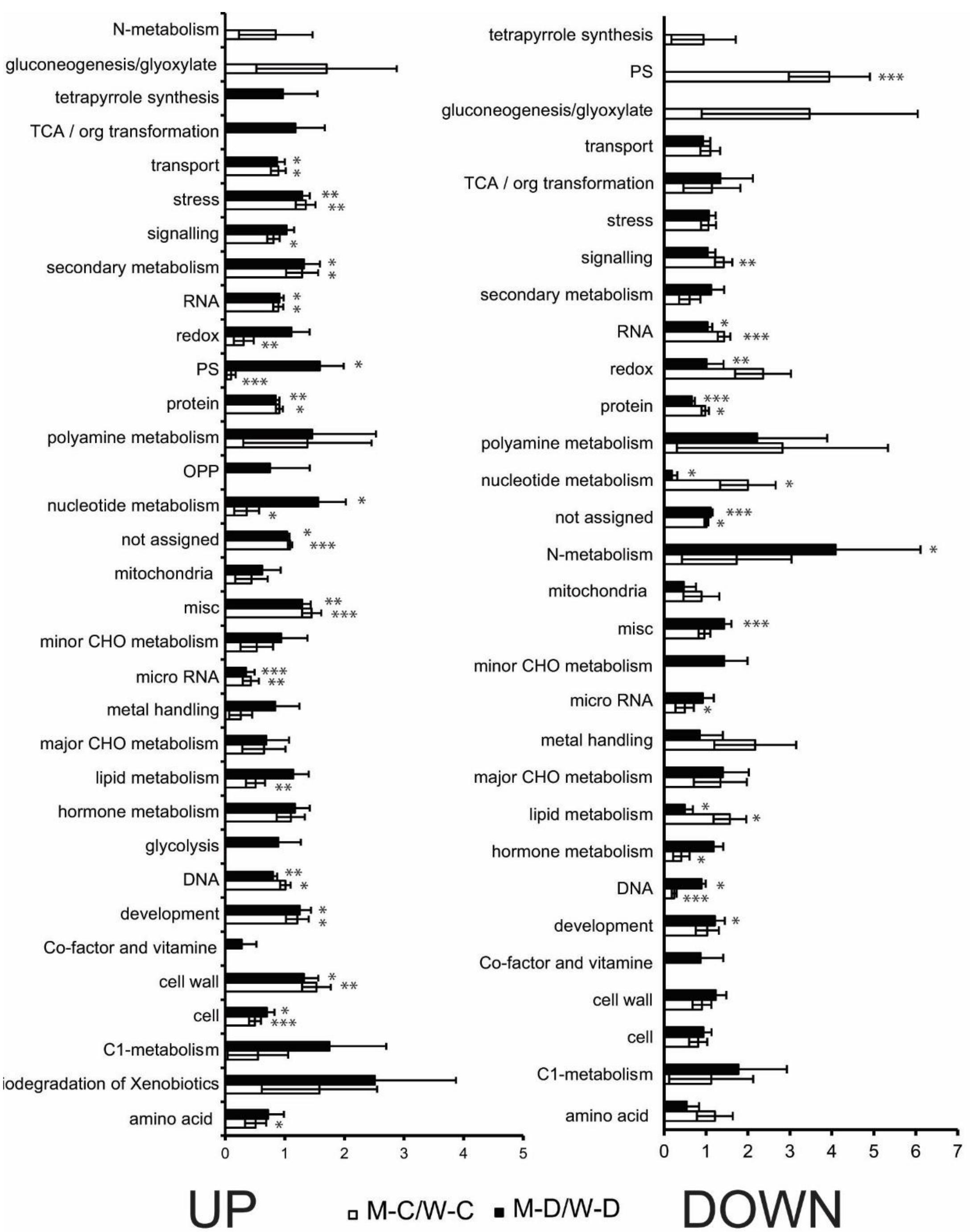


**Figure S4**. MapMan-based analysis of differentially expressed genes identified in *ahp4* mutant versus wild-type (WT) plants under normal (M-C/W-C comparison) and dehydration (M-D/W-D comparison) conditions. MapMan-based functional classification of the induced and repressed genes identified in M-C/W-C and M-D/W-D using the web-tool Classification Superviewer (bar.utoronto.ca) with normalized class score option (**P* < 0.05, ***P* < 0.01, ****P* < 0.001). CHO, carbohydrate; TCA, tricarboxylic acid; M-C/W-C comparison represents *ahp4* well-watered control versus WT well-watered control; M-D/W-D comparison represents M-D2/W-D2 and/or M-D5/W-D5; W-D2, WT dehydrated 2 h; W-D5, WT dehydrated 5 h; M-C, *ahp4* well-watered control; M-D2, *ahp4* dehydrated 2 h; M-D5, *ahp4* dehydrated 5 h.

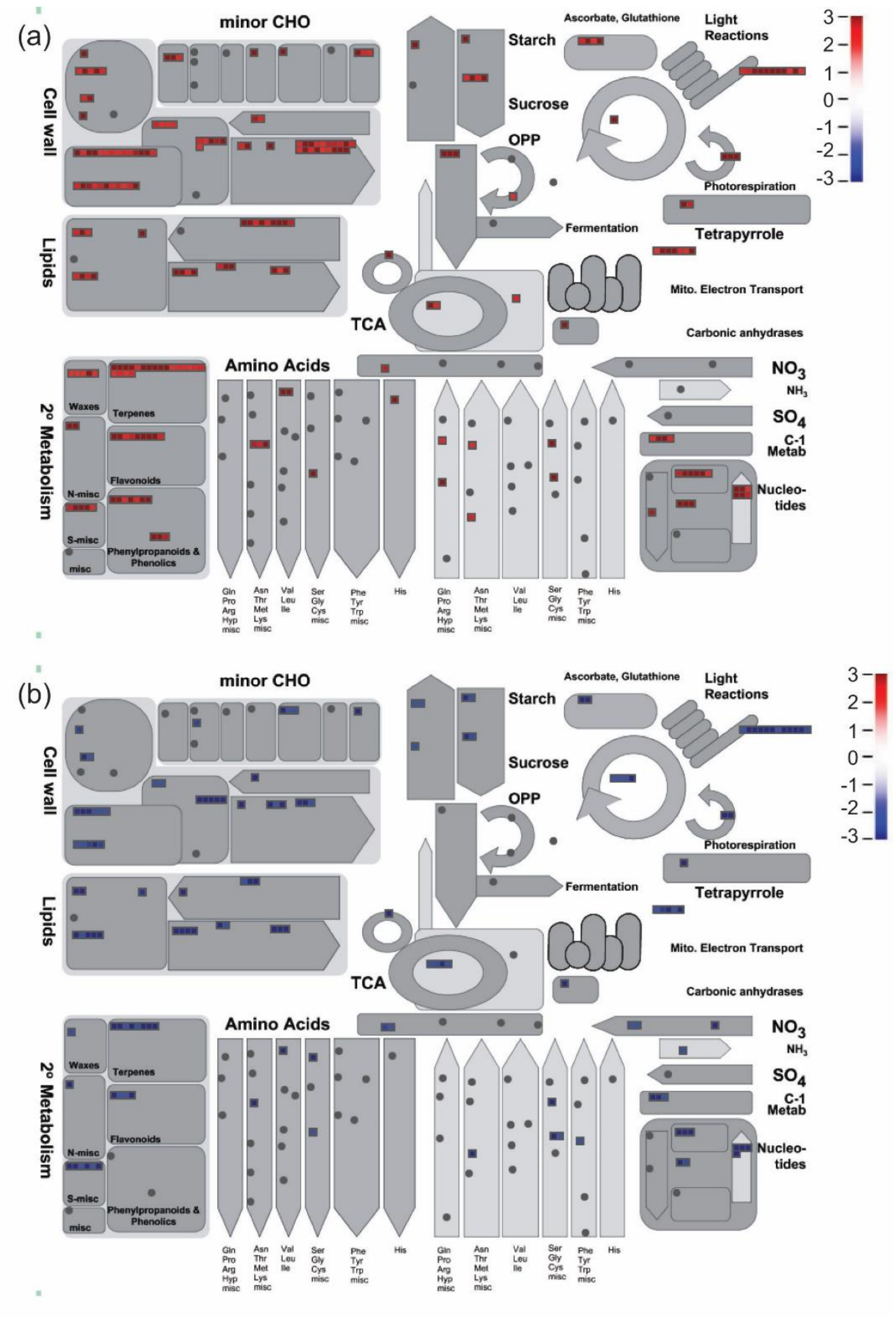


**Figure S5.** Metabolism-related overview of differentially expressed genes derived from *ahp4* versus wild-type under normal and dehydration conditions using MapMan. Metabolism-related overview of up-regulated (a) and down-regulated (b) genes. Color bar in each panel indicates fold changes in gene expression.

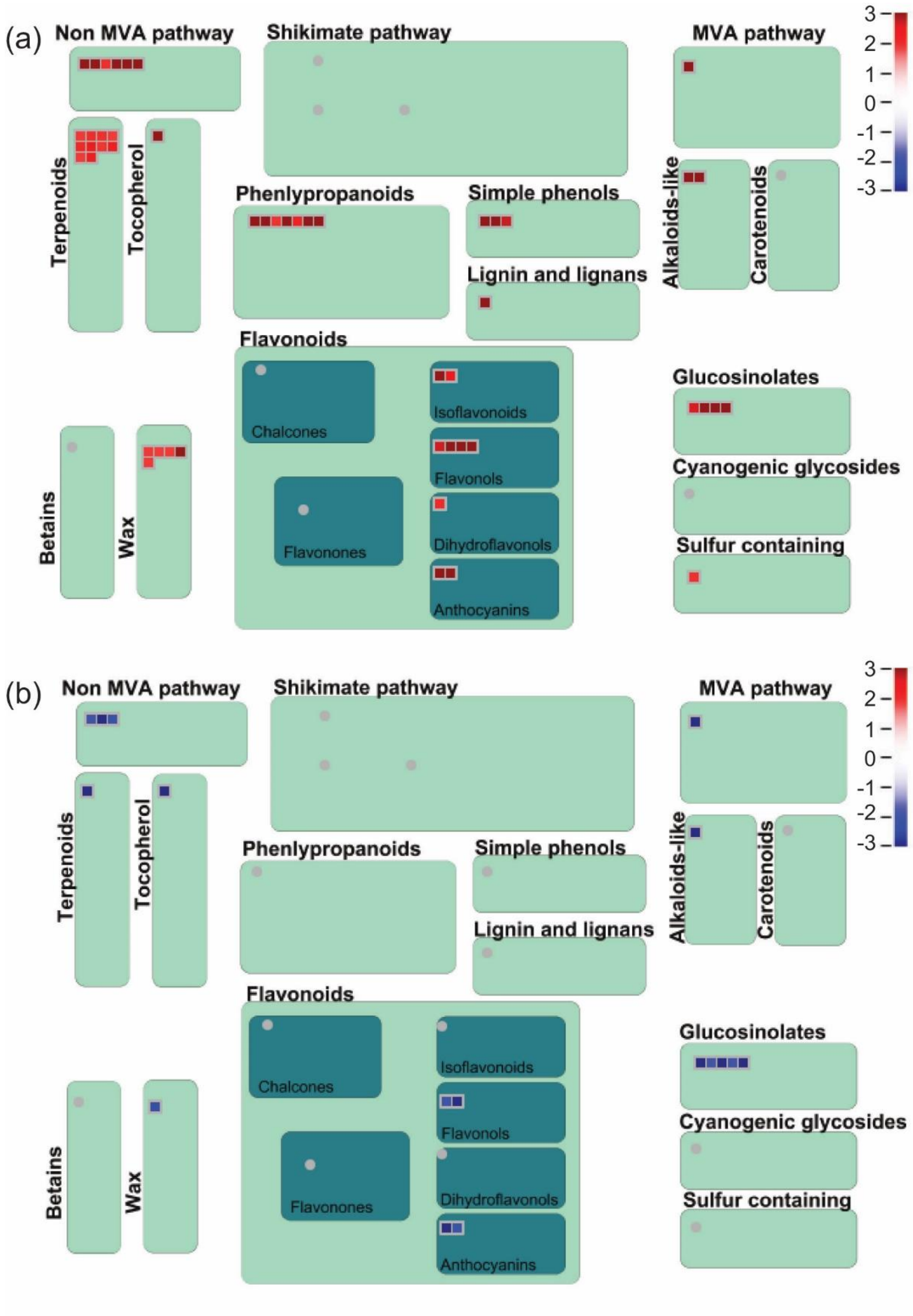


**Figure S6.** Secondary metabolism-related overview of differentially expressed genes identified in *ahp4* versus wild-type under normal and dehydration conditions using MapMan. (a) Up-regulated and (b) down-regulated genes. Red and blue colors indicate up- and down-regulated genes, respectively. Color bar in each panel indicates fold changes in gene expression.

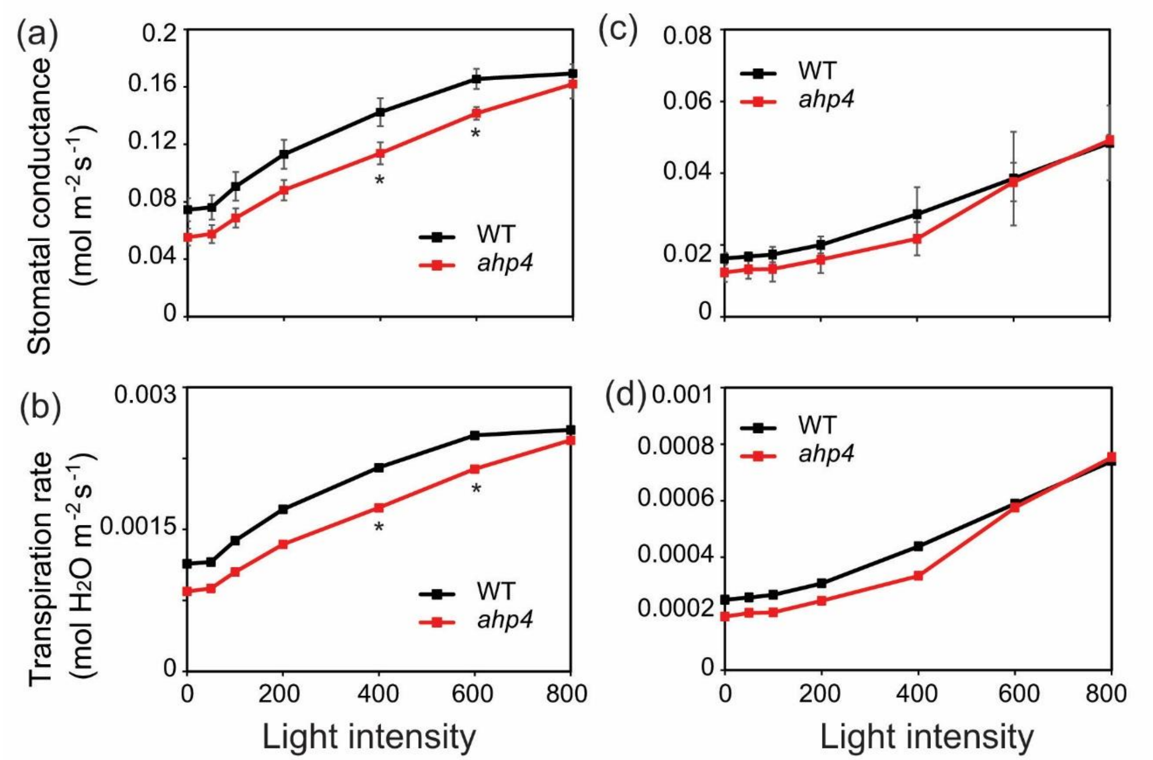


**Figure S7**. Stomatal conductance and leaf transpiration rates of *ahp4* mutant and wild-type (WT) plants under well-watered and dehydration conditions. (a) Stomatal conductance, and (b) leaf transpiration rates of 32-day-old *ahp4* and WT plants grown on soil under well-watered conditions. (c) Stomatal conductance, and (d) leaf transpiration rates of 32-day-old *ahp4* and WT plants grown on soil exposed to 30 minutes of dehydration treatment. Data represent the means and standard errors ($n = 3$). Asterisks indicate significant differences between the two genotypes as determined by a Student's $t$-test ($^*P < 0.05$).

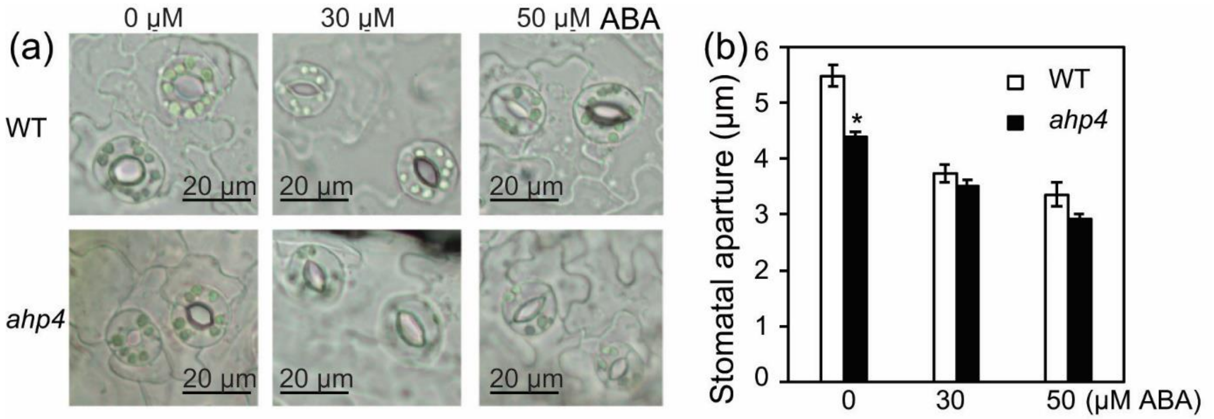


**Figure S8.** Comparison of stomatal apertures of *ahp4* mutant and wild-type (WT) plants under abscisic acid (ABA) treatment. (a) Guard cells of 21-day-old *ahp4* and WT plants were treated with 0 (control), 30 and 50 µM ABA for 1 h under light conditions. (b) Average size of the stomatal aperture of rosette leaves from 21-day-old *ahp4* and WT plants in the presence or absence of ABA. Data represent the means and standard errors ($n$ = 3 plants/genotype; for each plant the average of eight stomatal measurements from a single leaf was calculated). Asterisks indicate significant differences between the two genotypes as determined by a Student's $t$-test (*$P < 0.05$).

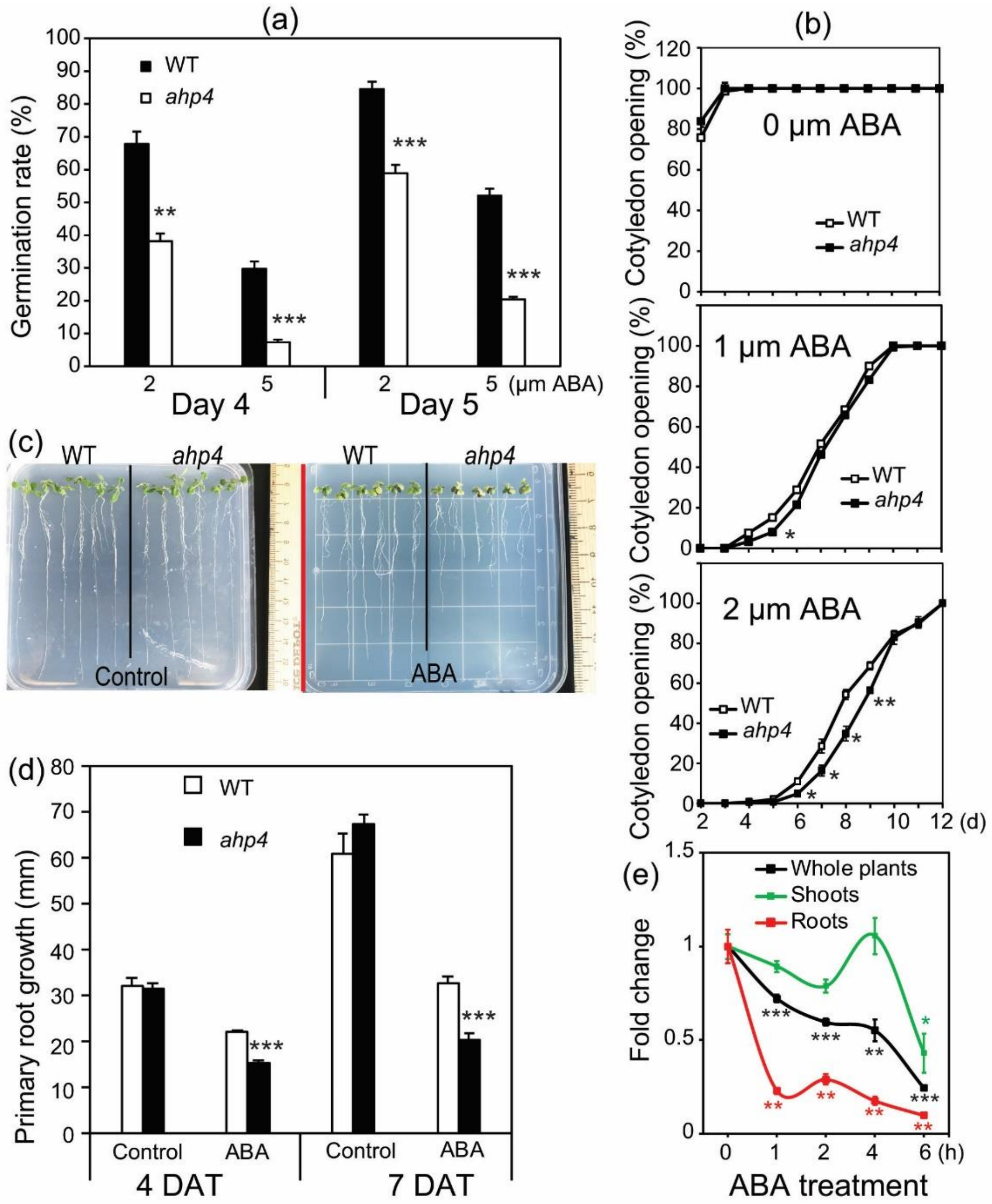


**Figure S9.** Germination assay of *ahp4* mutant and wild-type (WT) plants on medium supplemented with different concentrations of abscisic acid (ABA), and *AHP4* expression in WT plants treated with ABA. (a) Germination rate of *ahp4* mutant and WT plants under exogenous ABA treatment. Data represent the mean and standard errors (SEs) ($n$ = 8, where each replicate represents the radical emergence rate of ≥ 18 seeds/genotype). (b) Cotyledon opening percentage of *ahp4* mutant and WT under exogenous ABA treatment. Data represent the mean and SEs ($n$ = 3, where each replicate represents the cotyledon opening rate of 50 seeds/genotype). (c) Seven-day-old *ahp4* and WT plants grown on germination medium were transferred onto 0.5 × MS plates (1.2% w/v agar) with or without 20 µM ABA for 7 days. (d) Primary root growth of *ahp4* and WT plants under normal and ABA conditions as described in (c). Data present the mean and SEs ($n$ = 18 plants/genotype). (e) Expression of *AHP4* in 21-day-old WT plants treated with 50 µM ABA for indicated time period. Data represent the mean and SEs ($n$ = 3). Asterisks indicate significant differences as determined by Student's $t$-test (*$P < 0.05$, **$P < 0.01$, ***$P < 0.001$). DAT, days after transfer.

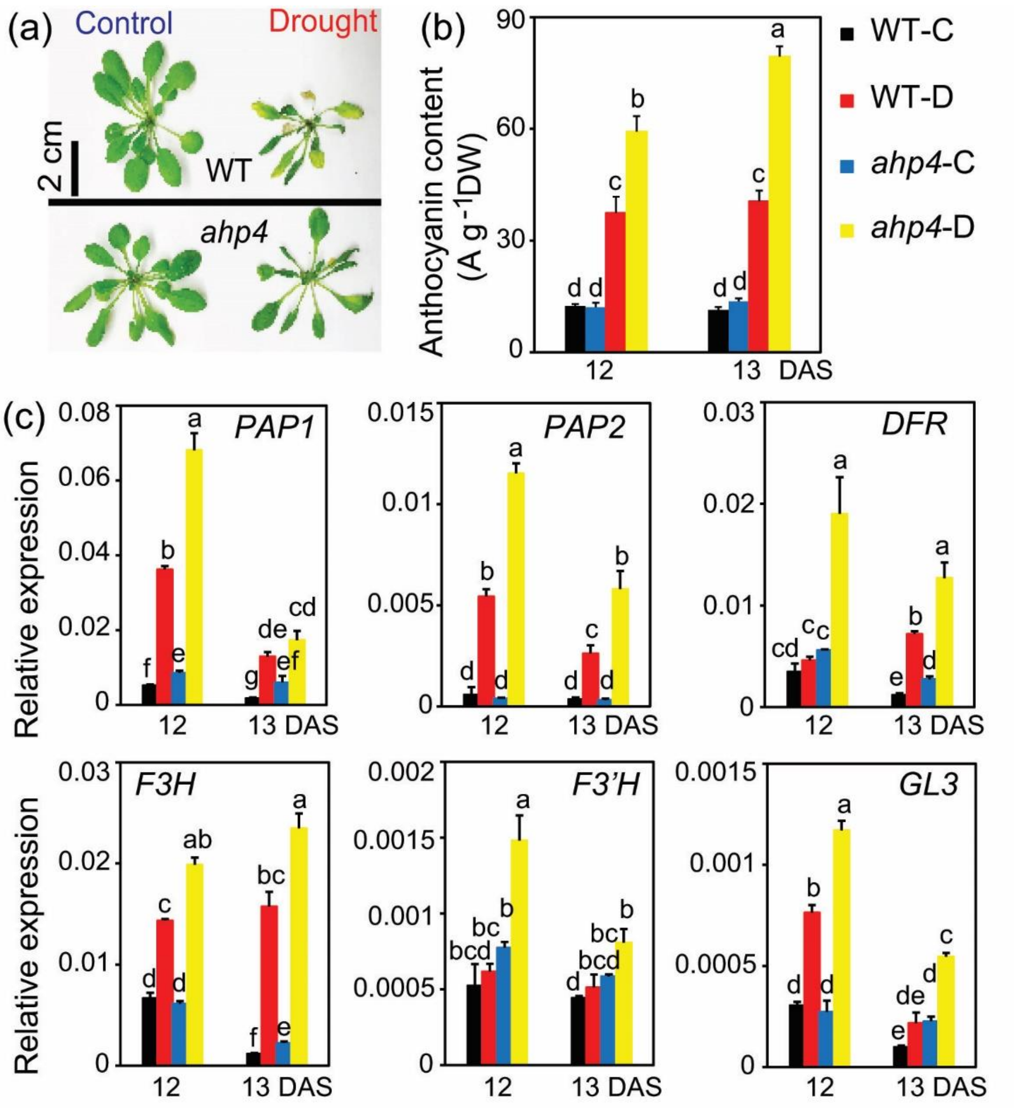


**Figure S10.** Anthocyanin contents and expression of anthocyanin/flavonoid-related genes in *ahp4* mutant and wild-type (WT) plants under drought. (a) Representative of *ahp4* and WT plants under control and 12 days after soil-drying. (b) Anthocyanin contents in *ahp4* and WT plants under control and drought conditions. (c) Expression of anthocyanin/flavonoid-related genes in *ahp4* and WT plants under control and drought conditions. Data represent the mean and standard errors ($n$ = 5). Different superscripted letters (a, b, c, d, e, f and g) within the column indicate statistically significant differences between the two genotypes, and among the treatments determined by Duncan's multiple range test ($P < 0.05$). DW, dry weight; DAS, days after stress; WT-C, wild-type control; WT-D, wild-type drought; *ahp4*-C, *ahp4* control; *ahp4*-D, *ahp4* drought; *PAP1*/*PAP2*, *production of anthocyanin pigment 1*/*2*; *DFR*, *dihydroflavonol 4-reductase*; *F3H*, *flavanone 3-hydroxylase*; *F3′H*, *flavonoid 3′-monooxygenase*; *GL3*, *glabra 3*.

**Table S1.** Results of the comparative microarray analysis of leaves of *ahp4* mutant and wild-type plants under well-watered and dehydration conditions.

**Table S2**. List of up- and down-regulated genes in M-C/W-C comparison (|fold-change| ≥ 2; q-value < 0.05). (a) List of up-regulated genes (fold change ≥ 2, q-value <0.05) in M-C/W-C comparison. (b) List of down-regulated genes (fold change ≤ - 2, q-value <0.05) in M-C/W-C comparison. M-C/W-C, *ahp4* well-watered control 0 h versus WT well-watered control 0 h.

**Table S3**. List of up- and down-regulated genes in various comparisons (|fold-change| ≥ 2; q-value < 0.05). (a) List of up-regulated genes (fold change ≥ 2, q-value <0.05) in M-D/W-D comparison. (b) List of up-regulated genes (fold change ≥ 2, q-value <0.05) in M-D2/W-D2 comparison. (c) List of up-regulated genes (fold change ≥ 2, q-value <0.05) in M-D5/W-D5 comparison. (d) List of down-regulated genes (fold change ≤ - 2, q-value <0.05) in M-D/W-D comparison. (e) List of down-regulated genes (fold change ≤ - 2, q-value <0.05) in M-D2/W-D2 comparison. (f) List of down-regulated genes (fold change ≤ - 2, q-value <0.05) in M-D5/W-D5 comparison. (g) List of up-regulated genes (fold change ≥ 2, q-value <0.05) in W-D/W-C comparison. (h) List of down-regulated genes (fold change ≤ - 2, q-value <0.05) in W-D/W-C comparison. M-D/W-D, *ahp4* dehydrated 2 and/or 5 h versus WT dehydrated 2 and/or 5 h; M-D2/W-D2, *ahp4* dehydrated 2 h versus WT dehydrated 2 h; M-D5/W-D5, *ahp4* dehydrated 5 h versus WT dehydrated 5 h; M-D/M-C, *ahp4* dehydrated 2 h and/or 5h versus *ahp4* well-watered control 0 h; W-D/W-C, WT dehydrated 2 h and/or 5h versus WT well-watered control 0 h..

**Table S4**. Venn analysis of differentially expressed gene sets derived from various comparisons. (a) List of up-regulated genes (fold change ≥ 2, q-value <0.05) in M-C/W-C and W-D/W-C. (b) List of up-regulated genes (fold change ≥ 2, q-value <0.05) in M-C/W-C and W-D/W-C. (c) List of down-regulated genes (fold change ≤ - 2, q-value <0.05) in M-C/W-C and W-D/W-C. (d) List of down-regulated genes (fold change ≤ - 2, q-value <0.05) in M-D/W-D and W-D/W-C. M-C/W-C, *ahp4* well-watered control 0 h versus WT well-watered control 0 h; W-D/W-C, WT dehydrated 2 h and/or 5h versus WT well-watered control 0 h; M-D/W-D, *ahp4* dehydrated 2 and/or 5 h versus WT dehydrated 2 and/or 5 h.

**Table S5**. List of photosynthesis-related genes in *ahp4* and wild-type (WT) leaves under normal and dehydration conditions.

**Table S6**. Primers used in RT-qPCR.
